# *Drosophila* primary microRNA-8 encodes a microRNA encoded peptide acting in parallel of *miR-8*

**DOI:** 10.1101/2021.03.22.434418

**Authors:** Audrey Montigny, Patrizia Tavormina, Carine Duboe, Hélène San Clémente, Marielle Aguilar, Philippe Valenti, Dominique Lauressergues, Jean-Philippe Combier, Serge Plaza

**Affiliations:** Université de Toulouse, UPS, CNRS, UMR5546, Laboratoire de Recherche en Sciences Végétales, 31320 Auzeville-Tolosan, France; Centre de Biologie du Développement (CBD), Centre de Biologie Intégrative (CBI), Université de Toulouse, CNRS, Bat 4R3, 118 route de Narbonne, F-31062, Toulouse, France

**Keywords:** *Drosophila*, sORF, lncRNA, *miR-8*, miPEP, small peptides

## Abstract

**Background:** Recent genome-wide studies of many species reveal the existence of a myriad of RNAs differing in size, coding potential and function. Among these are the long non-coding RNAs, some of them producing functional small peptides via the translation of short ORFs. It now appears that any kind of RNA presumably has a potential to encode small peptides. Accordingly, our team recently discovered that plant primary transcripts of microRNAs (*pri-miRNAs*) produce small regulatory peptides (miPEPs) involved in auto-regulatory feedback loops enhancing their cognate microRNA expression which in turn controls plant development. Here we investigate whether this regulatory feedback loop is present in *Drosophila melanogaster*.

**Results:** We perform a survey of ribosome profiling data and reveal that many pri-miRNAs exhibit ribosome translation marks. Focusing on miR-8, we show that *pri-miR-8* can produce a miPEP-8. Functional assays performed in Drosophila reveal that miPEP-8 affects development when overexpressed or knocked down. Combining genetic and molecular approaches as well as genome-wide transcriptomic analyses, we show that *miR-8* expression is independent of miPEP-8 activity and that miPEP-8 acts in parallel to *miR-8* to regulate the expression of hundreds of genes.

**Conclusion:** Taken together, these results reveal that several *Drosophila pri-miRNAs* exhibit translation potential. Contrasting with the mechanism described in plants, these data shed light on the function of yet un-described *pri-microRNA* encoded peptides in *Drosophila* and their regulatory potential on genome expression.

## Background

More than twenty years after the first genome annotation, it is now becoming clear that the protein-centric view of gene content strongly underestimates the number of DNA regions that are expressed and fulfil important functions for development and physiology since the majority of the genome is in fact transcribed [1]. A first discovery was the importance of hundreds of small non-coding RNAs, such as microRNAs (*miR*), playing regulatory roles in the silencing of genes and transposable elements. More recently, genome-wide transcript profiling has disclosed the existence of numerous RNAs referred to as long non-coding RNAs (lncRNAs or lincRNAs) since they lack the classical hallmarks of protein-coding genes.

Although the functions of all lncRNAs remain largely unknown, there are several experimental cases illustrating their key role as functional RNAs in various steps of the control of genome expression [2]. In association with other molecules, lncRNAs can coordinate several physiological processes and their dysfunction impacts several pathologies, including cancer and infectious diseases. lncRNAs control genetic information, such as chromosome structure modulation, transcription, splicing, messenger RNA (mRNA) stability, mRNA availability and post-translational modifications. They also act as scaffolds, bearing interaction domains for DNA, mRNAs, miRs and proteins, depending on both their primary sequence and secondary structure [3]. In addition, while lncRNAs annotated as non-coding cannot produce large-sized proteins, they all contain myriads of short open reading frames (sORF) [4–7] and a surprising result was the discovery for a subset of them of their translation into small functional peptides [8–11].

MicroRNAs define a class of small, non-coding RNAs able to down-regulate the expression of target mRNA by binding to the 3’-ends inducing mRNA degradation and/or translation repression. Intergenic microRNAs are produced from the sequential cleavage of long precursors named primary transcripts of microRNA (*pri-miRs*) (frequently annotated as lncRNAs) by Drosha and Dicer into 22nt miRNA duplexes associated with the RISC protein complex. Identified in a broad spectrum of living species, they are transcribed from coding genes or lncRNAs by the RNA polymerase II. MicroRNAs are critical for normal animal development and are involved in many biological processes [12]. Due to their role in silencing, *miRs*, and in particular the *pri-miR*s they come from, have always been considered as non-coding. However, the discovery that plant *pri-miR*s encode small regulatory peptides merely adds evidences that *pri-miRNAs* are also pre-mRNAs [13]. In plants, miPEPs specifically increase transcription of their primary transcript impacting the level of the mature miR produced and consequently affecting the control of the entire *miR* Gene Regulatory Network (GRN). To date, this regulation has been extended to several miRs in various plants [14–17]. In human cells, only few reports present evidences of pri-miR translation [18–21]. Pri-miR-22 host gene endogenously produces a miPEP for which the function is unknown [19]. miR-200 might produce a miPEP able to control the Vimentin, a miR200 target [18]. miPEP155 was described to control major histocompatibility complex class II-mediated antigen presentation by disrupting the HSC70-HSP90 machinery [20]. However, whether these miPEPs control the expression of their cognate miR was not investigated. More recently, miPEP133 was discovered in miR34a as a tumor suppressor localized in the mitochondria. It enhanced p53 transcriptional activation which, in turn, induces miR-34a expression [21]. In summary, whereas few human pri-miRNA appear translatable, it remains not clear whether the regulation found in plants exists in animals. Here we addressed this question in *Drosophila melanogaster*. We show that several intergenic *pri-miRs* contain marks of ribosome profiling. To investigate whether *Drosophila* can produce miPEPs, we focused on *miR-8*, a previously well-characterized microRNA that sustains many developmental traits [22]. *miR-8* controls organism physiology, tissue growth and survival [23–26], stem cell renewal [27–30], central nervous system development [30–32], signalling and developmental pathways [33–41]. Consequently, *miR-8* loss or gain of function impinges on fly development and survival. In the *pri-miR-8*, we located a small ORF encoding a potential 71 amino acid peptide we called miPEP-8. We showed that this sORF is translatable *in vitro* and *in vivo*. While our attempts to reveal an auto-regulatory loop remained unsuccessful, we showed by genetic and transcriptomic approaches that miPEP-8 and *miR-8* act in parallel in controlling wing development and in regulating the expression of distinct sets of genes.

## Results

### Translation potential of sORFs present in *Drosophila pri-microRNAs*

As plants *pri-miRs* contain sORFs (miORFs) producing functional miPEPs involved in a positive feedback loop on *pri-miR* expression (Figure 1A), we first asked whether *D. melanogaster pri-miRs* contain significant levels of miORFs. We scored the number of sORFs present in intergenic *miR* genes and their *pri-miR* and compared this number with other classes of RNAs, coding as well as non-coding (Figure 1B). *Pri-miRs* present the highest enrichment of sORFs/kb when compared with the 5’UTR of coding genes, sequences known to contain translatable short open reading frames, but contain similar amounts of sORFs when compared with lncRNAs, previously reported to be widely bound by ribosomes [42]. Ribosome profiling experiments were developed to study translatability of RNAs by scoring the sequences bound by ribosomes [7, 43, 44] including studies conducted on *Drosophila* [4, 45, 46]. We searched if and how many *Drosophila pri-miRs* were widely bound by ribosomes. Briefly, we searched in the rib-seq databases marks of ribosome binding in predicted *pri-miR*. To avoid difficulties for the interpretation with *miRs* embedded in host coding genes, we focused only on the intergenic *miRs*. We found many marks of ribosome profiling and identified hundreds of potentially translated sORF peptides within dozens of *pri-miRs*, suggesting that, as observed in plants, *Drosophila pri-miRs* are potentially translated (Additional file 1: Figure S1; Additional file 2).

**Figure 1:**
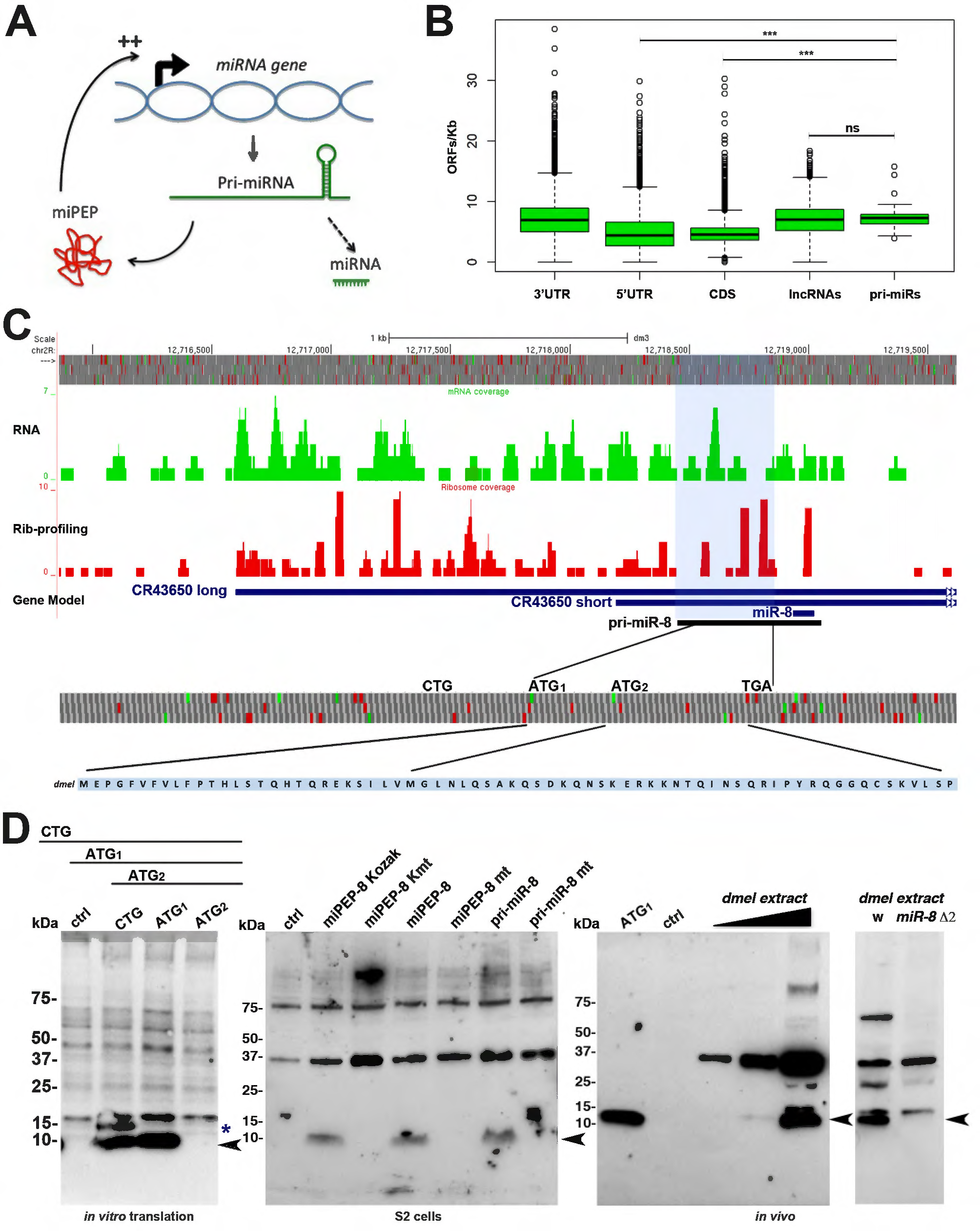
Translatability of *pri-miR-8*. **A:** Model of miPEP regulation in plants. **B:** Box plot representation of the number of ORFs in different classes of RNAs. 3’UTR, 5’UTR and CDS represent coding RNAs, whereas lncRNAs and pri-miRs represent non-coding RNAs. An ORF was defined as starting with an ATG and coding for a minimum of 10 amino acids. *Pri-miRs* reveal comparable numbers of ORFs/kb as lncRNAs. **C:** GWIPS-vis [65] genome viewer of the *Drosophila miR-8* locus. Top: genomic positions and ORFs in the three reading frames. Green bars define ATGs and red bars stop codons. Below, RNA-seq profile is shown in green and ribosome profiling is shown in red. The blue horizontal lines represent the two CR43650 non-coding RNA transcripts and potential *miR-8 pri-miRNAs*. In black is schematized the transcript we identified as detectable *pri-miR-8*. Bottom: miPEP-8 amino acid sequence is shown. **D:** western blot experiments using the anti-miPEP-8 antibody. Left panel: *in vitro* synthetized miPEP-8 corresponding to the constructs indicated on top. The asterisk indicates the upstream initiated peptide. The arrow indicates the miPEP-8 initiated at ATG1. Middle panel: detection of miPEP-8 in S2 cells over-expressing miPEP-8 placed in different translational contexts; Kozak (optimal); K mt (ATG mutated into TGA); mt (ATG mutated into AGT). Note that the *pri-miR* is translated and endogenous miPEP-8 expression is undetectable in S2 cells. Right panel: anti miPEP-8 western blot of adult *Drosophila* extracts and in the *miR-8* deleted line Δ2/Δ2 [26] in which no miPEP-8 is detected. We noticed the presence of non-specific bands as well as additional specific bands representing possibly miPEP-8 multimeric forms or PTM modifications. Ctrl corresponds to cell extracts transfected with an empty vector.

### *Drosophila pri-miR8* encodes a miPEP-8 translated *in vivo*

To further characterize the potential translation of *pri-miRs*, we focused on the *miR-8* primary transcript (*pri-miR-8). MiR-8* is an intergenic miR and is likely produced from the expression of long non-coding CR43650 spanning over the *pre-miR-8* (Figure 1C). In flybase, two CR43650 ncRNAs were predicted, a long and a short form defining putative *pri-miR-8* transcripts, independently identified by two different teams and likely initiated from different promoters [47, 48]. As shown in Figure 1C, many marks of potential translation were found along the CR43650 transcripts. We first performed 5’RACE as well as RT-PCR assays to determine which isoform was preferentially produced. While we successfully detected the short isoform in flies and S2 cells, we did not succeed in amplifying the long isoform, suggesting that the short transcript is the most abundant isoform expressed, a result confirmed by RNA-seq data generated during this study (Additional file 1: Figure S2). Our RNA-seq data further suggest that the long transcript might define a different transcription unit since no overlapping reads were detected in the promoter region of the short transcript (Additional file 1: Figure S3A). In agreement with this, a GAL4 enhancer trap recapitulating *miR-8* expression is inserted just upstream of the short transcrip*t* [24]. These flies express *miR-8* at a level comparable to control flies (Additional file 1: Figure S3B), showing that this insertion disrupting the co-linearity of the *miR-8* locus is not detrimental for *miR-8* expression. Finally, we verified that this short transcript (referred hereafter as *pri-miR-8*) is functional since it efficiently produces a functional mature *miR-8* able to down regulate a *miR-8* sensor (see below Figure 3B, D).

We therefore looked for a potential open reading frame within the *pri-miR-8* gene. Focusing on the 5’ leader sequence of *pri-miR-8*, we found one ORF located upstream of the *pre-miR-8*. This ORF is the longest ORF present 5’ to the *pre-miR*, which potentially encodes a miPEP of 71 amino acids in length if initiated from the first ATG (ATG1) (Figure 1C and Additional file 1: Figure S2A). However, the presence of a second ATG (ATG2), located downstream, gives the possibility to produce a shorter peptide. To determine whether the open reading frame is translated and which initiation codon is used, we generated and characterized specific antibodies (Additional file 1: Figure S4). In parallel, we generated different deletion constructs and performed *in vitro* translation experiments using insect cell extracts. As shown in Figure 1D (left panel), we observed an efficient translation from the longest construct (CTG) consisting in an extended genomic region of the defined 5’ leader sequence of *pri-miR-8*. Deletion experiments revealed a stronger and efficient translation from ATG1 but not from ATG2. A higher product, possibly initiated at an upstream codon present in the construct was also detected but was not further investigated since it was not present in *pri-miR-8*.

We next generated translatable and untranslatable miPEP-8 forms placed in optimal translational Kozak (K) or mutated kozak (KMT) contexts or in the miPEP-8 natural ATG1-initiated translational context. We then expressed these miPEP-8 constructs in *Drosophila* S2 cells (Figure 1D, middle panel). As revealed by western blot experiments, these different constructs produced the same level of miPEP-8 when the ATG was placed in an optimal or in its natural translational context whereas no miPEP-8 was detected from mutated ATG constructs. This result reveals that the natural nucleotide context of miPEP-8 miORF is in a favorable translational context. We next questioned whether *pri-miR-8* was able to produce miPEP-8. We transfected S2 cells with wild type or ATG-mutated *pri-miR-8* constructs and performed western blot experiments. We observed that only the wild-type *pri-miR-8* was able to produce miPEP-8. Finally, we examined whether a peptide corresponding to miPEP-8 was detectable in fly extracts by performing a western blot experiment on young adult flies. As revealed in Figure 1D (right panel), we observed a signal co-migrating with the *in vitro* synthetized miPEP-8, corresponding to endogenous miPEP-8 as demonstrated by the lack of this band in *miR-8* deleted Δ2 mutant flies. Sequence alignment analyses revealed that this peptide is poorly conserved amongst *Drosophila* species. Some homologies are detected in *Drosophila* melanogaster group but no conservation was found in more distant *Drosophilae* species.

Altogether, our results reveal that the *pri-miR-8* transcript carries, in addition to the *miR-8* sequence, at least one translated ORF located upstream of the *pre-miR*, able to express a miPEP-8 both *in vitro* and *in vivo*.

### Expression of miPEP-8 impinges on *Drosophila* development

To study the function of miPEP-8, we generated flies able to express a translatable and untranslatable version of miPEP-8 (ATG mutated). *miR-8* sustains many biological functions in *Drosophila* and either its loss of function or its over-expression leads to detrimental outcomes in cells, tissues or the whole organism [23–27, 29, 31–37, 40, 41, 49], providing a useful readout to assess miPEP-8 activity. To test our hypothesis that miPEP-8 controls *miR-8* expression and modulates its activity, we used an over-expression assay. Using the *miR-8* GAL4 driver, a GAL4 insertion in the endogenous *miR-8* promoter reported to mimic *miR-8* expression [23, 24, 33, 40], we first asked whether over-expression of miPEP-8 impinges on fly viability. As reported, driving UAS-*miR-8* over-expression results in increased fly lethality [23] (Figure 2A). Over-expression of a UAS-miPEP-8 translatable construct also affected fly viability whereas the untranslatable form did not. This indicates that, like *miR-8*, the translatable form of miPEP-8 is able to interfere with development, although with a weaker effect compared to *miR-8*.

**Figure 2:**
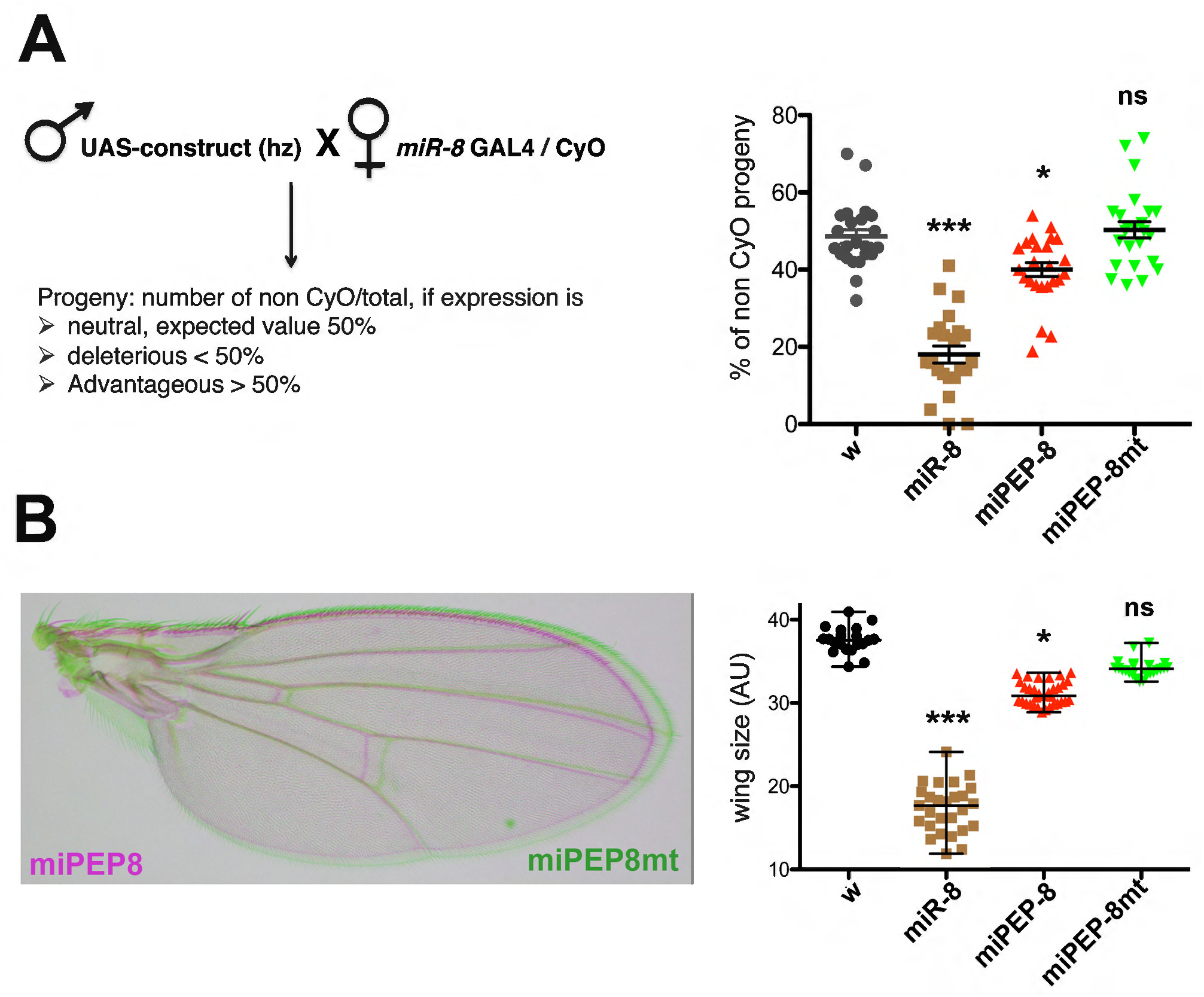
*Drosophila* miPEP-8 is biologically active during development. **A:** Lethality assay on flies over-expressing *miR-8*, miPEP-8 or miPEP-8mt (ATG mutated) using the *miR-8* GAL4 driver. Left: details of the genetic cross and expected percentage depending on the effect (neutral, deleterious or advantageous) on *Drosophila* development. Right: graph indicating the percentage of hatched flies over-expressing the different constructs. *white* flies (*w*) crossed with the driver line were used as a control. Expressing *miR-8* resulted in developmental lethality since less than 20% of flies hatched (expected value 50%). A significant decrease occurred following miPEP-8 over-expression but not with the untranslatable miPEP-8mt construct. Number of independent crosses: for *w* and *miR-8* n= 23; for miPEP-8 wt and mt n=24. **B:** Same as in **A** except the constructs were expressed in wings using the MS1096 driver and the phenotypes scored on wing size. Number of wings analyzed: for *w* n= 20; for *miR-8* n=27; for miPEP-8 wt and mt n= 27. * or ns: Significant differences are indicated relative to white recipient flies. AU: Arbitrary Units.

By loss or gain of function experiments, *miR-8* was shown to induce a « small wing » phenotype [24, 26, 40]. We therefore questioned whether over-expression of miPEP-8 also induced a wing phenotype (Figure 2B). Using the wing driver line MS1096, *miR-8* overexpression induced several wing defects, from a reduced size to the loss of wing vein, sensory organs, miss shaped, depending on the transgene/promoter strength [40, 41]. Quantifying the wing size appeared the most reliable criteria, and we compared this phenotype with the phenotype observed with miPEP-8 over-expression. Consistently with our above result, miPEP-8 over-expression induced a slight, albeit significant wing size reduction, revealing yet again a weaker activity compared with *miR-8* (Figure 2B). Importantly, miPEP-8-induced wing reduction was dependent on the integrity of the translation codon since the same construct with the mutated ATG did not induce any phenotype (Figure 2B).

Altogether, these experiments show that miPEP-8 appears to be biologically active but induces a milder phenotype compared to *miR-8*.

### In *Drosophila, miR-8* expression is independent of miPEP-8 expression

In plants, miPEPs positively auto-regulate the expression of their own *miR* by regulating the expression of their cognate *pri-miR*. To test whether miPEP-8 regulates *miR-8* expression in *Drosophila*, we monitored the level of *pri-miR-8* and *miR-8* through quantitative PCR experiments on S2 cells and on flies expressing the translatable form of miPEP-8. We first set up experimental conditions in S2 cells by transfecting the *pri-miR-8* construct and quantified the level of the exogenous *pri-miR* and mature miR produced. When *pri-miR-8* was over-expressed, both over-expression of *pri-miR-8* and *miR-8* was detectable (Figure 3B). We then over-expressed miPEP-8 and quantified the endogenous level of *pri-miR-8* and mature *miR-8*. Whereas we unambiguously detected the overexpression of the miPEP-8 construct, we did not see any change in the levels of endogenous *pri-miR-8* or mature *miR-8* expression (Figure 3C). As observed in S2 cells, no change in *pri-miR-8* or miR-8 levels was observed in flies upon miPEP-8 expression using the *miR-8* GAL4 driver (Additional file 1: Figure S5).

**Figure 3:**
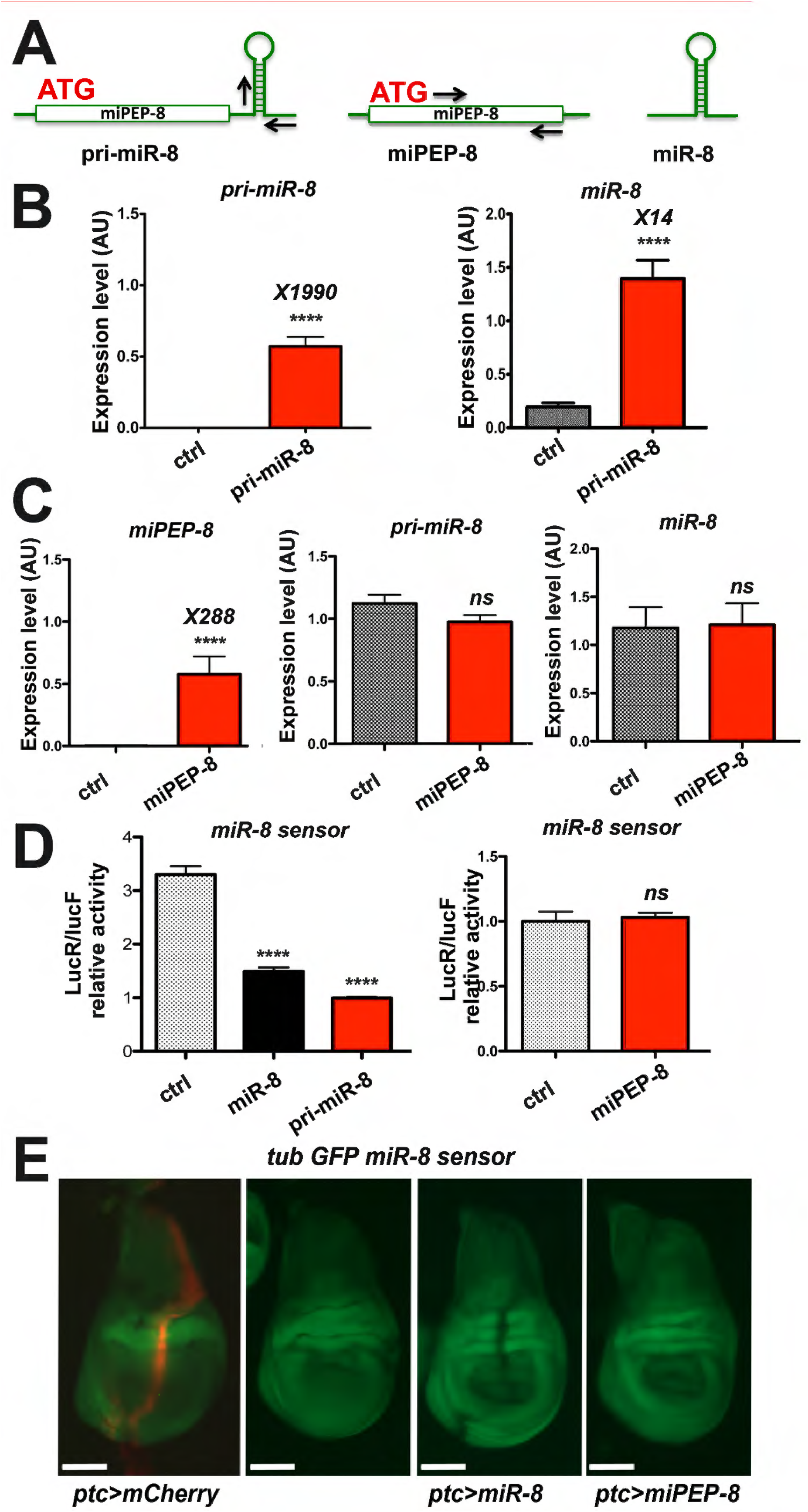
*Pri-miR-8* expression is independent of miPEP-8 control/activity. **A:** schematic representation of constructs tested on *miR-8* expression and activity levels. Arrows locate the primers used in the qPCR experiments determining miPEP and *pri-miR* relative expression levels. **B:** the characterized *pri-miR-8* produces a mature *miR-8*. S2 cells were transfected with a vector expressing *pri-miR-8*. Left: detection of the over-expression level of *pri-miR-8* by qPCR. Right: detection of the over-expression level of mature *miR-8* using the same RNA samples, n=11 **C:** miPEP-8 lacks repressive activity towards *miR-8* expression. Left: level of miPEP-8 over-expression. Middle and right panels: quantification of *pri-miR-8* and mature *miR-8* in miPEP-8 over-expressing cells compared to control transfected cells (ctrl). n= 13 for the ctrl and 14 for miPEP-8. **D** and **E**: Insensitivity of *miR-8* sensor to miPEP-8 over-expression in S2 transfected cells (n=16) (D) or in wing imaginal discs when miPEP-8 is expressed under the *ptc-GAL4* promoter (E). In D, a *miR-8* construct (n=12) [17] was used as a positive control repressing the *miR-8* luciferase sensor [20]. Of note, *pri-miR-8* (n=21) also repressed the *miR-8* luciferase sensor. In E, first panel to the left: *ptc* GAL4 crossed with a UAS mCherry. Second panel to the left: expression pattern of the GFP *miR-8* sensor alone. Scale bars (white) indicate 100mm. A repressive activity is observed with *miR-8* expressed in the *ptc* domain but not with miPEP-8. A representative disc is shown out of ten analysed.

We next questioned the potential regulatory role of miPEP-8 on *miR-8* expression by testing miPEP-8 overexpression on endogenous miR-8 activity level in the presence or absence of. One way of challenging this question is the use of a sensor of miR-8 activity, whether endogenous or resulting from over-expression. Thus, we designed a miR-8 luciferase reporter, bearing a 3’UTR from the *escargot* gene (*esg*), previously shown directly regulated by *miR-8* [27]. Over-expression of *pri-miR-8* in S2 cells was able to repress the miR-8 sensor to the same extent as *miR-8* (Figure 3D, left panel), hence validating our miR-8 sensor. As mentioned above, this also indicates that the *pri-miR-8* construct is able to generate a functional and mature *miR-8*. Clearly, however, over-expression of miPEP-8 did not reveal any modulation of the luciferase reporter (Figure 3D right panel). We performed similar experiments *in vivo* using a miR-8 GFP sensor in wing imaginal discs where *miR-8* was previously shown to be functional [33]. Expressing *miR-8* under the patched (ptc) GAL4 promoter led to the repression of the GFP in the ptc domain (Figure 3E ptc>miR8 panel). Consistently, miPEP-8 over-expression had no effect on the miR-8 GFP sensor in vivo (Figure 3E ptc>miPEP-8 panel). Altogether, these results indicate that miPEP-8 is not able to control *miR-8* expression, or activity for that matter (see more below). We obtained similar conclusions on endogenous miR-8 target in S2 cells and in wing discs (Additional file 1: Figure S6).

Finally, we asked whether the miPEP-regulation of *pri-miR* observed in plants is system specific by testing whether a plant *pri-miR*, up regulated by its miPEP in plant cells, could be up-regulated in *Drosophila* cells. Reciprocally, we tested whether miPEP-8 is able to up-regulate its *pri-miR-8* in plant cells. To that end, we expressed the plant *Arabidopsis thaliana pri-miR165a* and miPEP-165a in S2 cells using the actin promoter and measured the level of *pri-miR* produced in the absence or presence of the miPEP. Reciprocally, the *Drosophila pri-miR-8* and its miPEP-8 were cloned in plant expression vectors and agroinfiltrated in *Nicotiana benthamiana* leaves as performed previously [13]. Whereas the up-regulation of the *A. thaliana pri-miR-165a* by miPEP-165a was observed in *N. benthamiana*, we did not detect any up-regulation, but rather a down-regulation of *pri-miR-165a* in *Drosophila* S2 cells (Additional file 1: Figure S7). Reciprocally, we could detect a slight but significant increase of *pri-miR-8* expression upon miPEP-8 over-expression in *N. bentamiana* leaves, suggesting that a difference of regulation occurs between plant cells and insect cells (Additional file 1: Figure S7). Although, *miR-8* expression appears to be miPEP-8 independent in *Drosophila*, these results further suggest that, like for plants miPEPs, animal miPEPs might nonetheless have the potential of autoregulating the expression of their cognate pri-miR.

### Endogenous miPEP-8 alteration reveals *in vivo* activity

To investigate the functional requirement of miPEP-8 *in Drosophila*, we tried several times to edit the miPEP-8 in flies using CRISPR/Cas9, but unsuccessfully. In contrast, it was possible from the first attempt to delete the entire *miR-8* locus, showing that the failure to obtain a specific miPEP-8 edited line is not due to trivial technical problems. We therefore created a specific P landing platform in place of *pri-miR-8* transcript to perform Knock In strategies (Figure 4A and Additional file 1: Figure S8). This edited line exhibits the previously *miR-8* reported phenotypes [26], including a strong developmental lethality with only few escaping flies exhibiting a reduced size (including wings (Figure 4C and Additional file 1: Figure S8)) and leg defects (not shown). We further knocked in the wild type *pri-miR-8* and the *pri-miR-8* miPEP-8 untranslatable form (mt) at the P landing site and analyzed the outcomes. For both constructs, we observed a nearly total rescue since the theoretical expected 33,3 % homozygotes (and 66,6 % of CyO flies) in the progeny was almost reached (Figure 4B left panel). These rescued flies appeared phenotypically normal and re-expressed *miR-8* at levels close to *miR-8* endogenous expression (Figure 4B right panel). Both pri-miR-8 constructs restored significantly wing sizes when compared to the Δ*miR-8* CRISPR line (Figure 4C). Interestingly, a significant difference was observed in wings between the *pri-miR-8* wild type construct and the *pri-miR-8 mt* in which the miPEP-8 translatability was disrupted (Figure 4C).

**Figure 4:**
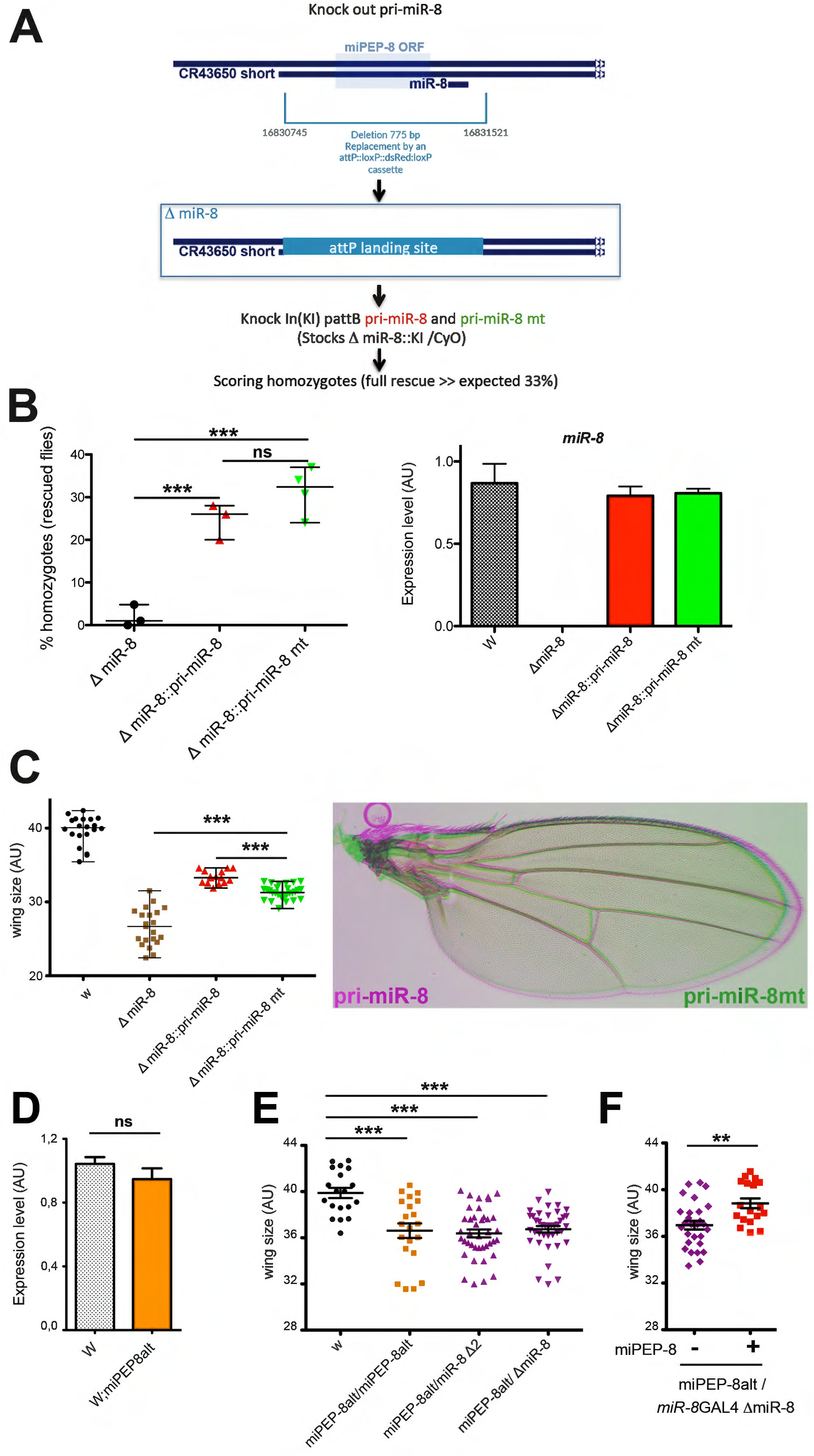
targeting miPEP-8 *in vivo* in *Drosophila* induces a wing phenotype. **A:** Strategy for endogenous miPEP-8 edition. The *pri-miR-8* gene region was deleted by CRISPR and a P landing site was created. Wild type and miPEP-8 ATG mutated *pri-miR-8* in pattB were inserted at the P landing site. **B**: Similar rescue efficiency was observed in at least three independent transgenic lines (left panel). qPCR on mature miR-8 in wild type and mutant (mt) *pri-miR-8* Knock In (KI) lines showed similar miR-8 levels (n=4). (right panel) **C**: wing phenotype in *miR-8* deletion edited line. The *pri-miR-8* miPEP-8 mutated (mt) shows a reduced wing size compared to the wild type *pri-miR-8*. (n= 15 and 28 respectively). D to F: analyses in miPEP-8 mutant identified in DGRP polymorphism **D**: miR-8 level determined by qPCR in white recipient flies (w) and in white flies carrying the miPEP-8 truncated form (miPEP-8alt). (n=6 and 8 respectively) **E** and **F**: Wing size determination in different genetic contexts. miPEP-8alt homozygotes or over *miR-8* deficiencies revealed significant reduced wing size relative to the white recipient flies (w, n=19; miPEP-8alt, n= 21; miPEP-8alt/ miR-8 deltions, n=40). Expressing miPEP-8 rescued the wing phenotype of miPEP-8alt flies relative to sibling flies not expressing miPEP-8 (n= 18 and 28 respectively). Significant (*) or non significant (ns) differences are indicated either relative to white recipient flies or between the two groups.

As a second approach, we took advantage of a polymorphism mutation detected in *Drosophila* Gene Reference Panel (DGRP) lines generating a premature stop codon leading to a 24 amino acid C-terminal miPEP-8 truncation called miPEP-8alt [50]. We outcrossed the miPEP-8alt DGRP line into white background and analyzed the consequence of the miPEP-8alt mutation. Whereas no significant difference was observed for miR-8 level between these the two miPEPs variants (Figure 4D), flies homozygous for miPEP-8alt exhibit a significant wing size reduction when compared with the white flies expressing the miPEP-8 (Figure 4E). We further analyzed the resulting wing phenotypes in different genetic contexts. The phenotype is also present when miPEP-8alt mutation was tested over a deletion of the *miR-8* gene (the Δ2 and the Δ*miR-8* CRISPR line generated in this study) (Figure 4E), suggesting that the observed phenotype is a consequence of miPEP-8 loss of function. To test this, we performed rescue experiments by expressing miPEP-8 using the *miR-8* GAL4 driver line in miPEP-8alt/Δ*miR-8* background (Figure 4F). Flies expressing miPEP-8 in miPEP-8alt/ Δ*miR-8* CRISPR restored the wing size phenotype contrasting with the sibling control flies carrying no miPEP-8 transgene (absence of expression of wild type miPEP-8) (Figure 4F).

Therefore, altogether, these experiments revealed an *in vivo* miPEP-8 function.

### The function of miPEP-8 is uncoupled from *miR-8* expression and activity

The above experiments suggest that in *Drosophila* miPEP-8 is not involved in a positive auto-regulatory feedback loop as observed in plants. However, due to the similarities of the phenotypes observed between *miR-8* and miPEP-8, we questioned whether miPEP-8 could be involved in the *miR-8* pathway through another mechanism, or whether it acts in parallel of miR-8. As both *miR-8* and miPEP-8 affected wing formation, we developed a genetic assay to test whether miPEP-8 acts through miR-8 using a previously validated miR-8 sponge, which titrates *miR-8* hence rescuing *miR-8*-induced phenotypes [29, 33]. Using this rescue assay, we asked whether the miR-8 sponge could also compensate the miPEP-8 induced phenotype. Co-expressing *miR-8* together with a miR sponge scramble (as a control and to maintain the number of UAS transgenes identical) using the MS1096 GAL4 driver led to wing size reduction. This phenotype was efficiently rescued by co-expressing *miR-8* with the effective miR-8 sponge (Figure 5). In contrast, when miPEP-8 was coexpressed with the *miR-8* sponge, no compensation of the miPEP-8-induced wing reduction was observed. Therefore, this result strongly suggests that miPEP-8 acts in parallel of *miR-8*.

**Figure 5:**
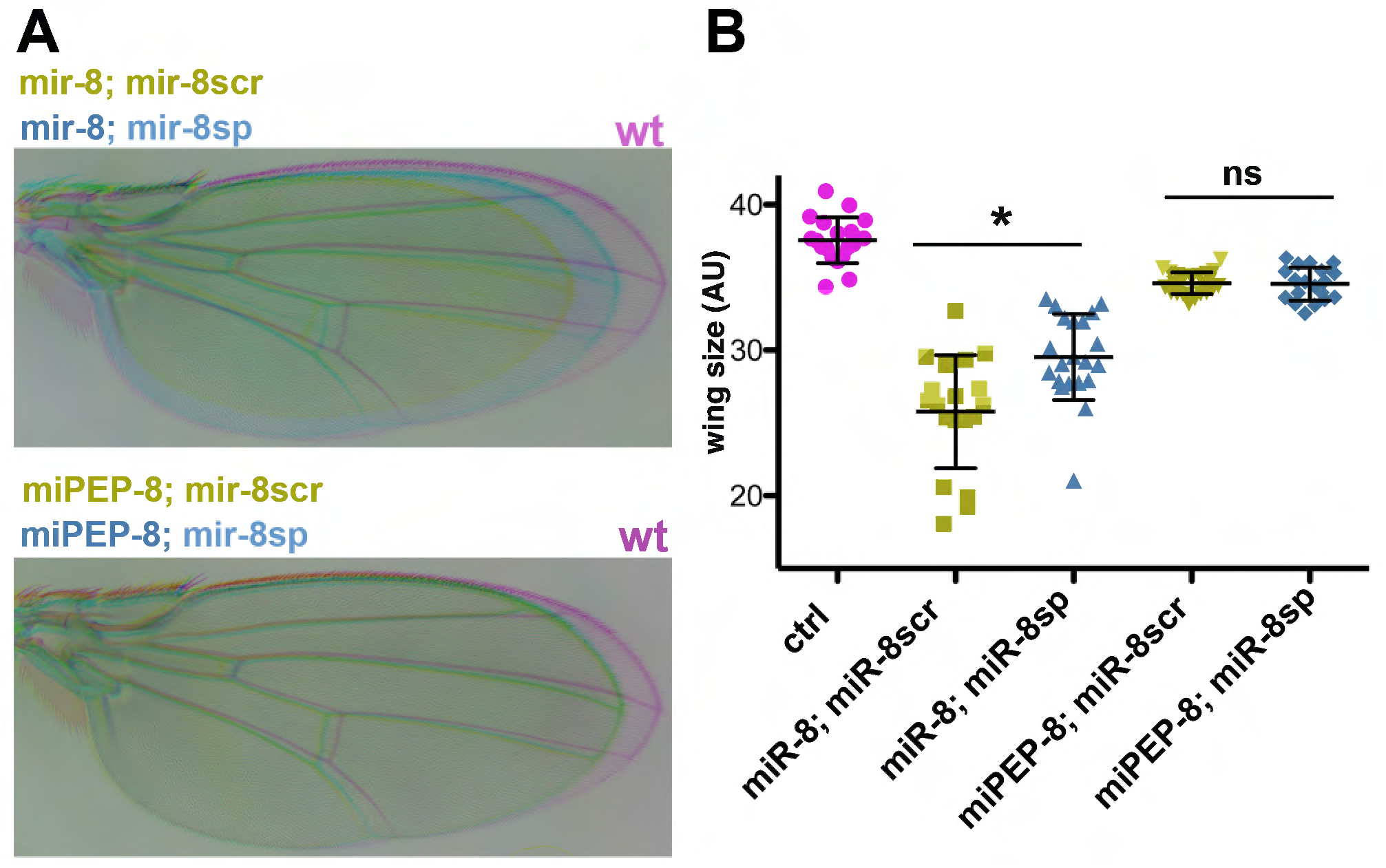
Uncoupled activity of miR-8 and miPEP-8. **A**: Rescue assay of *miR-8*- or miPEP-8- induced wing phenotype in flies co-over-expressing *miR-8* or miPEP-8 along with a miR-8 sponge (miR-8sp) or a miR-8 scramble (miR-8scr). Only miR-8-sp (and not miR-8scr) compensates for *miR-8*-induced wing size reduction, hence efficiently titrating *miR-8*, while it has no effect on miPEP-8-induced wing phenotype. **B**: Quantification of A. “ctrl” (MS1096/+) n=19; “mir-8; mir-8scr” n=20; “mir-8; mir-8sp” n=21; “miPEP-8; mir-8scr” n=23; “miPEP-8; mir-8sp” n=19. * p<0,05.

We thus reasoned that the effect of miPEP-8 on wing development could be linked to the modulation of gene expression independent of *miR-8*. To identify these putative miPEP-8-regulated genes and to compare them with the *miR-8*-regulated transcriptome, we overexpressed *miR-8* or miPEP-8 in S2 cells and performed RNA-seq 48h after transfection (Figure 5; Additional file 3). Clearly, the transcriptomes appeared different (Figure 6A; Additional file 1: Figure S9). The assays performed on *miR-8* over-expressing cells successfully retrieved previously identified *miR-8* targets both at the RNA (Additional file 1: Figure S9A and Additional file 2) and protein level (Additional file 1: Figure S6), hence validating our experimental conditions. GO term enrichment identified biological pathways fitting with miR-8 activity such as “regulation of organism or cell growth and differentiation”, “wing development”, “apoptosis”, “regulation of actin cytoskeleton” (Additional file 1: Figure S10). As for miPEP-8 controlled genes, strikingly, the majority of them were miPEP-8 specific (76%) (Figure 6B and 6C) since only 24% appeared co-regulated (Figure 6B and 6E). In both cases, we found activated and repressed genes (Figure 6A, C, D, E). Remarkably, miPEP-8-modulated genes were frequently more strongly modulated than *miR-8*-modulated genes (Figure 6C, D). Increasing the Fold change (FC>1,5) led to a decrease of the number of genes but the respective proportions and conclusions remained unchanged (Additional file 1: Figure S9B). Our analyses of GO term enrichment clearly identified shared functions for *miR-8* and miPEP-8 (Additional file 1: Figure S10A and S11A; Additional file 4), some of which being related to wing morphogenesis (such as cell junction organization actin filament-based processes, epithelial cell morphogenesis, cell differentiation) or developmental processes (such as neurogenesis, cell migration, embryonic morphogenesis) (Additional file 4). However, *miR-8* and miPEP-8 also exhibit specific biological functions such as snRNA modification and leucine metabolic process for miR-8 or K48 linked ubiquitination and chromatin-mediated maintenance of transcription for miPEP-8 (Additional file 1: Figure S10B and S11B).

**Figure 6:**
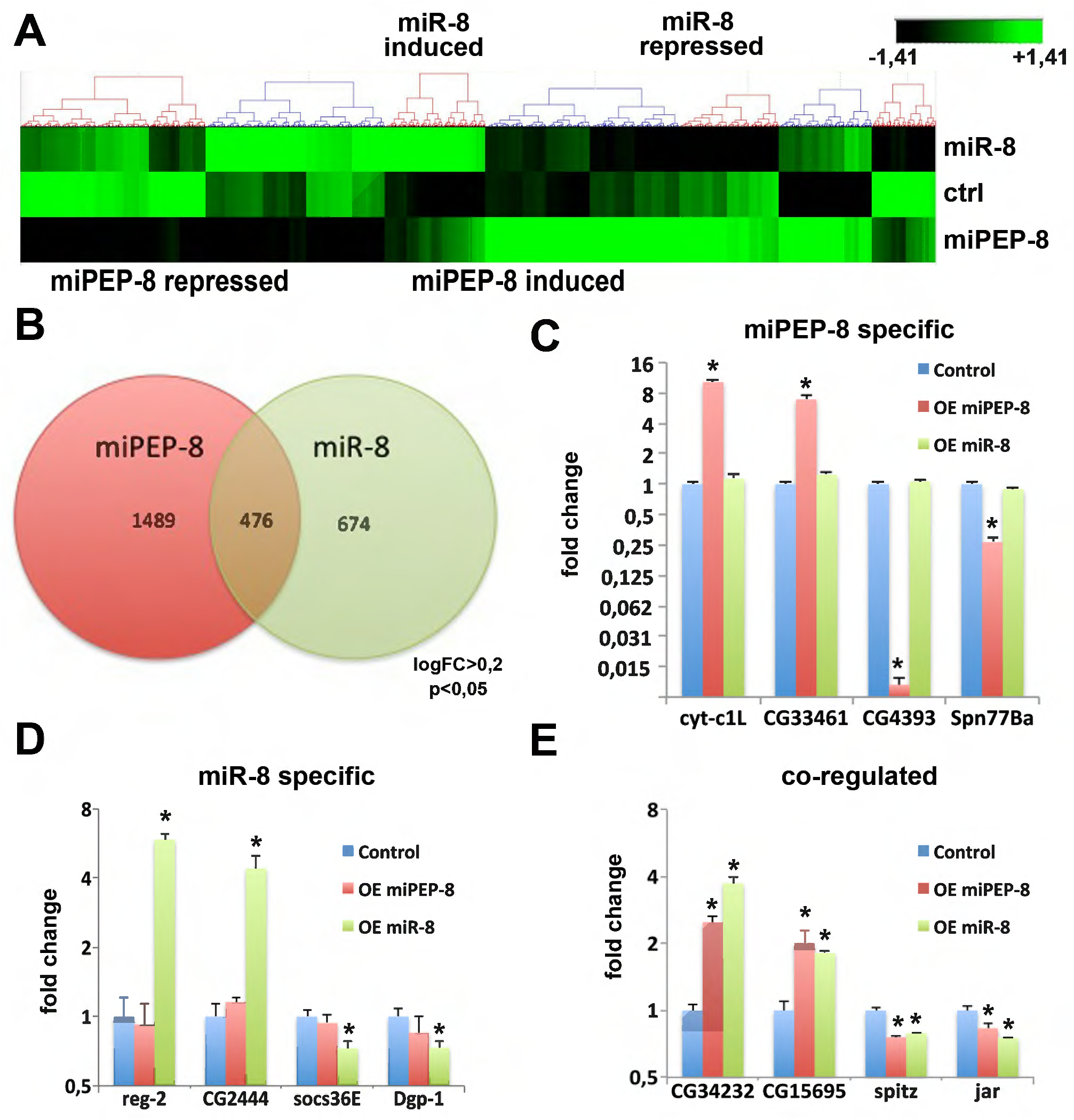
miR-8 and miPEP-8 control distinct set of genes. **A**: Heatmap representing the RNA-seq results obtained from S2 cells over-expressing either *miR-8* or miPEP-8. Significant sets of genes are modulated in response to *mirR-8* or miPEP-8 over-expression, when compared to control transfected cells (ctrl). N=5. **B:** Venn diagram representing the *miR-8* versus miPEP-8 modulated genes. **C, D, E**: different subgroups are distinguished; miPEP-8 specific (**C**), *miR-8* specific (**D**), and co-regulated by miPEP-8 and *miR-8* (**E**).

Altogether, these experiments suggest that *miR-8* and miPEP-8 independently control similar biological processes, while regulating functions specific to one or the other.

## Discussion

In the present study, we investigated whether a small ORF present in *Drosophila pri-miR-8* was capable of producing a miPEP-8 and we propose that animal miPEPs are able to act in parallel of their corresponding miRs. Several studies performed in a broad range of organisms have revealed the prevalence of translated small/short open reading frames (smORFs/sORFs) [7, 43, 44, 51–56]. Although sORF peptides were initially identified as being encoded by unusual long non-coding RNAs, to date, it turns out that many classes of RNAs can produce these peptides. Therefore, sORF-encoded peptides (SEPs) are emerging as an unexplored reservoir of putative regulators. However, while a growing body of evidence further supports the importance of sORFs and associated peptides in development, physiology and diseases [8, 54, 57, 58], the number of SEPs that have been characterized so far still remains limited. Therefore, the current challenge resides in deciphering the full repertoire of their functions and molecular modes of action, an issue largely dependent on experimental approaches.

We show with several experimental data that a miPEP-8 is indeed produced from *Drosophila pri-miR-8*. First, we found a signal of ribosome binding in the *pri-miR* of several microRNAs and in particular, *miR-8*. Second, we show that the initiation codon of the miORF present within *pri-miR-8* is in a favorable translational context. Third, after having generated specific antibodies, we detected a peptide co-migrating with *in vitro* translated miPEP-8 in fly extracts. Fourth, forced expression or loss of function of this peptide led to a significant developmental phenotype in *Drosophila* and induced significant variations of cellular gene expression. Therefore, the poor conservation detected amongst *Drosophila* species indicates that this sORF-encoded peptide differs from the few conserved ones characterized so far and shed light on it by its recent invention.

Here, we tackled the question of whether the miPEP auto-regulatory function was identical to that of plants. While we did not detect any auto-regulatory loop (miPEP increasing the expression of its own *pri-miR* and miR), we observed that the action of miPEP-8 is uncoupled from *miR-8* regulation. On the one hand, our data suggest that this peptide could control similar developmental outcomes or developmental pathways and share the regulation of identical subsets of genes. In this context, we further analyzed whether we could detect a significant miPEP-8 activity in other *miR-8* developmental processes such as intestinal stem cell differentiation [27] and eye morphogenesis [37] (data not shown). However, no significant activity was detected, suggesting that, in the experimental conditions tested, miPEP-8 does not act in all *miR-8* developmental pathways. Such an example was observed in S2 cells in which *miR-8* is expressed at detectable levels whereas endogenous miPEP-8 is not. On the other hand, we reveal that miPEP-8 likely has a regulatory function all of its own, independently of *miR-8*. Indeed, our RNA-seq data indicates that miPEP-8 regulates specific genes and biological processes (i.e. independent of *miR-8* activity). This also occurs *in vivo* since few candidates of the top list of miPEP-8 specific regulated genes identified in S2 cells are also modulated in miPEP-8 loss of function in adult flies (Additional file 1: Figure S12). Future loss of function and expression pattern analyses throughout development should bring further insight into miPEP-8-specific regulatory functions.

Is the uncoupling of miPEP activity from miR regulation a general feature of animal miR genes? The study performed here suggests that the mechanisms involved in animals might be different from the miPEP auto-regulatory mechanism observed in plants. As such, a recent study on human miR155 revealed an activity for a miPEP155 that is not correlated to miR155 control [20] and on human miR34 where a miPEP133 mitochondrial function impinging on p53 activity was reported [21]. In light of these results, of course, we cannot affirm that the mechanisms described here are common to all *Drosophila miR* genes. It remains possible that some of them might be auto-regulated by their miPEPs as described in plants. In addition, since ribosome occupancy were not found in all *Drosophila* microRNA genes, it remains possible that some *pri-miRs* are unable to produce miPEPs. Therefore, additional studies will be required to determine whether miPEP-dependent *pri-miR* autoregulation is specific or widespread amongst *miR* genes.

Is the *pri-miR* coding capacity conserved throughout the animal kingdom? In a search for non-coding RNAs able to express sORF-encoded peptides, Razooki and co-workers found that human *miR-22* host gene (*pri-miR-22*) produces a potential miPEP-22 that is induced during viral infection [19]. sORFs have also recently been identified in *miR-200a* and *miR-200b pri-miRs*, the human orthologs of the *Drosophila miR-8*. Like *miR-200a* and *miR-200b*, miPEP-200a and b over-expression in prostate cancer cells inhibits migration of these cells by regulating the vimentin-mediated pathway, suggesting that the miPEP-coding function of *pri-miRs* is present in humans [18]. Accordingly, most recently, micropeptides encoded by MIR155HG and MIR34HG were described to be involved in autoimmune inflammation by controlling antigen presentation [20] and mitochondrial function respectively [21] via their interaction with different HSP proteins. It is interesting to note however, that these miPEPs appear to be involved in infections/pathologic-conditions, hence suggesting that revealing miPEP function might be largely dependent on the biological context.

Ribosome-associated lncRNAs has been considered to constitute a hallmark of protein translation. Here we found a signal of ribosome binding in the *pri-miR* of several Drosophila *microRNAs* genes. Furthermore, we showed that the initiation codon of the miORF present within *pri-mR-8* is in a favorable translational context. Indeed, after having generated specific antibodies, we detected a peptide co-migrating with *in vitro* translated miPEP-8 in fly extracts. However, an alternative possibility proposed by others is that ribosome marks illustrate a mechanism for cellular control of lncRNA levels through ribosome degradation-promoting activity [56, 59]. It will be of interest to investigate further whether the short ORFs present in *pri-miRs* are able to influence their regulation by controlling their stability and degradation as it has been shown for other coding genes. Finally, the molecules that give rise to miR-8/miPEP-8 are probably not the same ones since Drosha processing would separate the ORF from the poly(A) tail and thereby cause rapid decapping and degradation of the ORF-containing fragment. In light with these considerations, it is difficult to conclude on a pervasive coding capacity of *pri-miRs* in *Drosophila*. Future work will determine both in plants and animals whether all of them are sources of miPEPs and to what extent their auto-regulatory capacity and/or modes of action are diverse and specific.

### Conclusion

Many studies performed recently have led to functional characterization of a handful of additional SEPs in the plant and animal kingdom. Illustrating the diversity of functions of these new players, these SEPs were identified from different sources of RNAs and play different roles [9, 60, 61]. Among these, contrasting with their initial definition as non-coding, pioneer works in plants showed that even precursors transcript of miRNAs produces SEPs involved in an autoregulatory feedback loop. By addressing the conservation of this mechanism in animals, our findings combined with others confirm that miR-encoded genes probably represent evolutionary conserved bi-functional RNAs carrying coding and non-coding functions. However, contrasting with the mechanism described in plants, our data shed light on the diverse functions fulfilled by microRNA-encoded-peptides despite their poor conservation among *Drosophila species*.

## Methods

### Fly strains and Genetics

*Drosophila* flies were maintained on standard cornmeal-yeast medium (Dutscher). Experiments were performed at 25°C when *miR-8* GAL4 (NP5247) was used as driver. For the experiments of wing phenotype of flies expressing transgenes under the control of MS1096 Gal4, crosses were placed at 28°C. UAS-*pri-miR*-8 and UAS-miPEP-8 transgenic lines were inserted in attP86F site through PhiC31-mediated integration. Injections were performed by Bestgene Inc (USA). Generating pri-miR-8 fly founder line: pri-miR-8 fly founder line was designed and generated by inDroso Functional Genomics (Rennes, France) using CRISPR/Cas9. The pri-miR-8 fly founder line was generated by excising from position 16830745 to 16831521 on Chromosome 2R arm and replacing it by an attP::loxP::3xP3-dsRED::loxP cassette (Additional file 1: Figure S8). The two following guide RNA sequences were used to cut on either side of the *pri-miR-8:* CACATATG|CAACGGAAAGAG and GTTGGTGG|TACTGAAGGTTA. The edited region was verified by sequencing. The two pri-miR-8 constructs in pattB were inserted at the Δ*miR-8* created P site. Three independent transformants were used for analyses. The same strategy was used to generate the driver *miR-8* GAL4, Δ*miR-8*. The miPEP-8 alternative form creating a premature stop codon in miPEP-8 was derived from the DGRP-859 line, outcrossed into white recipient flies and kept over CyO. Experiments are the sum of at least 3 independent crosses. n indicates the number of individuals analysed. For wing measurements, young flies (2-5 days) of the appropriate genotypes were stored in Ethanol. For analysis of wings, females adult wings were removed in wash buffer (PBS and 0.1% Triton X-100), and mounted on a slide in 80% glycerol in PBS as described [62]. Wings or wing discs images were acquired on a Zeiss Axiozoom stereomicroscope. Measurements of wing size were performed using IMAGE J software.

### Molecular methods

For cloning procedure, miPEP-8, miR-8 or pri-miR-8 plasmids were constructed from PCR amplification of genomic DNA, gene synthesis or by RNA reverse transcription from S2 cells or adult *Drosophila* RNA and cloned in pUAS-attB vector constructs using the In-fusion HD cloning kit (Takarabio) according to the manufacturer specification. All constructs were verified by sequencing. For quantitative PCR experiments, total RNA was isolated from young adult fruit flies (2-5 days) or S2 cells using TRI Reagent (Sigma) according to the manufacturer specifications, followed by RQ1DNase treatment (promega) according the manufacturer specifications. The cDNA template was synthesized using SuperScript III (Invitrogen) with oligo-dT18 as anchor primers. Quantitative real-time PCR was performed on the LightCycler 480 Instument II (Roche Life Science) using LightCycler480 SYBR GREEN I master (Roche Life Science). The mRNA abundance of the examined genes was estimated by qPCR. For the endogenous *pri-miR* or coding genes, RP49 and tubulin genes were used as reference genes and used for normalization. For quantifying mature miRNA, stem loop PCR conditions were set up and the small RNAs U14 and Sno442 were used as reference. Datas presented are the same whatever the reference gene used. When the S2 cells are transiently transfected, the co-transfected pActin-GAL4 vector (Addgene # 24344) was used to monitor transfection efficiency. Typically, each experiment presented was performed with four replicates processed independently and was repeated in time at least three times. All experiments were taken into account and “n” indicate the total number of biological replicates used for analyses. Primers used in the qPCR are listed in the Additional File 1: Figure S13.

### RNA analysis

For Rib-seq analyses, Dmel primiR three frames translations have been performed with transeq (Emboss suite 6.6.0). A homemade script written in Perl was generated to compare the resulting translated peptides to the ribo-seq sORF encoded peptides described in [4, 45, 46]. RNA-seq was processed by genewiz (Germany). Each dataset contains five independent biological replicates of control *miR-8* and miPEP-8 over-expressing S2 cells RNA-seq. The reads were subjected to standard quality control (QC) and filtered according to the following parameters: (1) trimming and cleaning reads that aligned to primers and/or adaptors, (2) reads with over 50% of low-quality bases (quality value ≤15) in one read, and (3) reads with over 10% unknown bases (N bases). We used Trimmomatic software (v0.36) to remove primers and bad quality reads. After filtering, we removed short reads (parameters were used with default values. Gene and PSI lists for each dataset were compared to identify common events between them. For RNAseq analysis, htseq-counts files were analyzed using the version 3.24.3 of package EdgeR [63], in order to normalize raw counts by “trimmed mean of M-values” (TMM), and test differential expression using the negative binomial distribution. RNA-seq analysis: raw p-values were adjusted with the Benjamini–Hochberg procedure to control the False Discovery Rate (FDR). A gene was declared differentially expressed if it’s adjusted p value ≤ 0.05. Heat map parameters applied: row-by-row normalisation by standardisation (Mean and Standard deviation). GO term analysis was performed with PANTHER (http://pantherdb.org/)[64]. Sashimi plots were created with IGV (Integrative Genomics Viewer, https://igv.org/)[65]. Statistical analyses were performed using the version 3.5.2 of R software and Bioconductor packages. For QPCR analysis, the version 1.3-1 of package Agricolae was used.

### Cell culture and western blot and luciferase assays

*Drosophila* S2 cells were maintained in Schneider’s medium (Invitrogen) supplemented with 10% fetal bovine serum (Sigma), 50 U/ml penicillin and 50 μg/ml streptomycin (Invitrogen) at 25°C. For western blot experiments, miPEPs sequences cloned into pF25A ICE T7 Flexi vector were expressed in vitro using TnT^®^ T7 Insect Cell Extract Protein Expression System (Promega). For cells extracts and *Drosophila* extracts, we directly freeze them in nitrogen just before western blot. Proteins were prepared in Laemli buffer (63 mM Tris HCl pH7.5, 2% SDS, 5% 2-mercaptoethanol) and run on SDS-PAGE according to [13]. Primary antibodies used for western were: rabbit anti-miPEP-8 were raised against the sequence KQSDKQNSKERKKNTQI (generated and affinity purified by Agro-bio, France), mouse anti-GAPDH (ThermoFisher AM4300), rabbit anti-Sra-1 (1/1000, provided by A. Giangrande, IGBMC CNRS, France), mouse anti-peanut (1/100, DSHB, USA) and rabbit anti-ABP-1 (1:250, provided by Michael Kessels Jena University Hospital, Germany). HRP conjugated secondary antibodies are from Santa Cruz Biotechnology (1/10000 sc-516102). For luciferase assays, in each experiment, S2 cells were transfected in quadruplicate, in 24-wells plates (700000 cells/well) using FuGene HD transfection Reagent (Promega). Experiments were repeated timely independently at least 3 times. After 48h of transfection, cells were washed with PBS and lysed with 100μL Passive Lysis Buffer (Dual luciferase Reporter Assay System, Promega). Firefly luciferase (FL) and Renilla luciferase (RL) activities were then quantified with DUAL luciferase reporter assay (Promega) using 50μl of reagents/well and measure using a Greiner luminometer instrument.

### Statistical analyses

Statistical analyses were performed using GraphPad Prism and illustrated as follow: * p-value<0,05; ** p-value<0,01; *** p-value<0,001; **** p-value<0,0001. In all experiments, results represent mean ±s.e.m. n represente the number of biological independent replicates. Normality test were first performed using D’Agostino Pearson test. If the distribution is Gaussian and in order to detect a global difference between all groups, oneway ANOVA was performed using one-way analyses of variances followed by Bartlett’s test for equal variance and Bonferroni’s multiple comparison tests. In other cases, when variance or sample sizes are not equal, non-parametric analyses were performed using Kruskal-Wallis test to detect a global difference between all groups followed by comparisons between two groups performed using adjustments for multiple comparisons. When only two groups were compared, a Mann & Witney test was performed.

## Declarations

### • Ethics approval and consent to participate

Not applicable

### • Consent for publication

Not applicable

### • Availability of Data and Materials

The RNAseq datasets generated during the current study are available in the NCBI Gene Expression Omnibus (GEO) repository under the accession number BioProject ID PRJNA645280 [66] [https://www.ncbi.nlm.nih.gov/Traces/study/?acc=PRJNA645280&o=acc_s%3Aa]

### • Authors’ contributions

SP and JPC conceived the project. SP designed the research. AM, PT, CD, PV, DL SP performed the experiments. HSC and MA performed bioinformatics analyzes. SP, AM, PT, JPC wrote the paper.

## • Acknowledgements

We thank C.Dozier and the « Peptides & small RNAs » team members for critical reading and their helpful comments on the manuscript. We thank H.Boukhatmi, A.Giangrande, S.Cohen, F.Juge for sharing flies, constructs, and antibodies. We thank the FRAIB RIO Imaging facility for microscopy. We acknowledge the Bloomington *Drosophila* Stock Center (NIHP40OD018537) and the Drosophila Stock Center, KYOTO Stock Center (DGRC) in Kyoto Institute of Technology for providing flies stocks, the Developmental Studies Hybridoma Bank (DSHB) for antibodies, and DNASU and Addgene for plasmids.

---

**Additional File 1:** Supplementary Figures S1 to S13

**Additional File 2:** list of sORF peptides defined from Rib-seq analyses

**Additional File 3:** normalized RNA-seq values from S2 cells.

**Additional File 4:** Protein ANalysis THrough Evolutionary Relationships (PANTHER) Overrepresentation Test.

## References

1. Cech TR, Steitz JA: The noncoding RNA revolution-trashing old rules to forge new ones. Cell 2014, 157:77–94.

2. Gardini A, Shiekhattar R: The many faces of long noncoding RNAs. FEBS J 2015, 282:1647–1657.

3. Ransohoff JD, Wei Y, Khavari PA: The functions and unique features of long intergenic non-coding RNA. Nat Rev Mol Cell Biol 2018, 19:143–157.

4. Aspden JL, Eyre-Walker YC, Phillips RJ, Amin U, Mumtaz MA, Brocard M, Couso JP: Extensive translation of small Open Reading Frames revealed by Poly-Ribo-Seq. Elife 2014, 3:e03528.

5. Bazzini AA, Johnstone TG, Christiano R, Mackowiak SD, Obermayer B, Fleming ES, Vejnar CE, Lee MT, Rajewsky N, Walther TC, Giraldez AJ: Identification of small ORFs in vertebrates using ribosome footprinting and evolutionary conservation. EMBO J 2014, 33:981–993.

6. de Andres-Pablo A, Morillon A, Wery M: LncRNAs, lost in translation or licence to regulate? Current Genetics 2017, 63:29–33.

7. Ingolia NT, Lareau LF, Weissman JS: Ribosome profiling of mouse embryonic stem cells reveals the complexity and dynamics of mammalian proteomes. Cell 2011, 147:789–802.

8. Plaza: In search of lost small peptides. Annual reviews 2017.

9. Rathore A, Martinez TF, Chu Q, Saghatelian A: Small, but mighty? Searching for human microproteins and their potential for understanding health and disease. Expert Rev Proteomics 2018, 15:963–965.

10. Ruiz-Orera J, Alba MM: Translation of Small Open Reading Frames: Roles in Regulation and Evolutionary Innovation. Trends Genet 2019, 35:186–198.

11. Yeasmin F, Yada T, Akimitsu N: Micropeptides Encoded in Transcripts Previously Identified as Long Noncoding RNAs: A New Chapter in Transcriptomics and Proteomics. Front Genet 2018, 9:144.

12. O’Brien J, Hayder H, Zayed Y, Peng C: Overview of MicroRNA Biogenesis, Mechanisms of Actions, and Circulation. Front Endocrinol (Lausanne) 2018, 9:402.

13. Lauressergues D, Couzigou JM, Clemente HS, Martinez Y, Dunand C, Becard G, Combier JP: Primary transcripts of microRNAs encode regulatory peptides. Nature 2015, 520:90–93.

14. Couzigou JM, Andre O, Guillotin B, Alexandre M, Combier JP: Use of microRNA-encoded peptide miPEP172c to stimulate nodulation in soybean. New Phytol 2016, 211:379–381.

15. Sharma S, Badola PK, Bhatia C, Sharma D, Trivedi PK: miRNA-encoded peptide, miPEP858, regulates plant growth and development in Arabidopsis. BioXiv 2019, 642561.

16. Chen QJ, Deng BH, Gao J, Zhao ZY, Chen ZL, Song SR, Wang L, Zhao LP, Xu WP, Zhang CX, Ma C, Wang SP: An miRNA-encoded small peptide, vvi-miPEP171d1, regulates adventitious root formation. Plant Physiol 2020.

17. Zhang QL, Su LY, Zhang ST, Xu XP, Chen XH, Li X, Jiang MQ, Huang SQ, Chen YK, Zhang ZH, Lai ZX, Lin YL: Analyses of microRNA166 gene structure, expression, and function during the early stage of somatic embryogenesis in Dimocarpus longan Lour. Plant Physiol Biochem 2020, 147:205–214.

18. Fang J, Morsalin S, Rao V, Reddy ES: Decoding of Non-Coding DNA and Non-Coding RNA: Pri-Micro RNA-Encoded Novel Peptides Regulate Migration of Cancer Cells. J Pharm Sci Pharmacol 2017, 3:23–27.

19. Razooky BS, Obermayer B, O’May JB, Tarakhovsky A: Viral Infection Identifies Micropeptides Differentially Regulated in smORF-Containing lncRNAs. Genes (Basel) 2017, 8.

20. Niu LM, Lou FZ, Sun Y, Sun LB, Cai XJ, Liu ZY, Zhou H, Wang H, Wang ZK, Bai J, Yin QQ, Zhang JX, Chen LJ, Peng DH, Xu ZY, Gao YY, Tang SB, Fan L, Wang HL: A micropeptide encoded by lncRNA MIR155HG suppresses autoimmune inflammation via modulating antigen presentation. Science Advances 2020, 6.

21. Kang M, Tang B, Li JX, Zhou ZY, Liu K, Wang RS, Jiang ZY, Bi FF, Patrick D, Kim D, Mitra AK, Yang-Hartwich Y: Identification of miPEP133 as a novel tumor-suppressor microprotein encoded by miR-34a pri-miRNA. Molecular Cancer 2020, 19.

22. Sander M, Herranz H: MicroRNAs in Drosophila Cancer Models. Adv Exp Med Biol 2019, 1167:157–173.

23. Jin H, Kim VN, Hyun S: Conserved microRNA miR-8 controls body size in response to steroid signaling in Drosophila. Genes Dev 2012, 26:1427–1432.

24. Karres JS, Hilgers V, Carrera I, Treisman J, Cohen SM: The conserved microRNA miR-8 tunes atrophin levels to prevent neurodegeneration in Drosophila. Cell 2007, 131:136–145.

25. Umegawachi T, Yoshida H, Koshida H, Yamada M, Ohkawa Y, Sato T, Suyama M, Krause HM, Yamaguchi M: Control of tissue size and development by a regulatory element in the yorkie 3’UTR. Am J Cancer Res 2017, 7:673–687.

26. Hyun S, Lee JH, Jin H, Nam J, Namkoong B, Lee G, Chung J, Kim VN: Conserved MicroRNA miR-8/miR-200 and its target USH/FOG2 control growth by regulating PI3K. Cell 2009, 139:1096–1108.

27. Antonello ZA, Reiff T, Ballesta-Illan E, Dominguez M: Robust intestinal homeostasis relies on cellular plasticity in enteroblasts mediated by miR-8-Escargot switch. EMBO J 2015, 34:2025–2041.

28. Antonello ZA, Reiff T, Dominguez M: Mesenchymal to epithelial transition during tissue homeostasis and regeneration: Patching up the Drosophila midgut epithelium. Fly (Austin) 2015, 9:132–137.

29. Boukhatmi H, Bray S: A population of adult satellite-like cells in Drosophila is maintained through a switch in RNA-isoforms. Elife 2018, 7.

30. Morante J, Vallejo DM, Desplan C, Dominguez M: Conserved miR-8/miR-200 defines a glial niche that controls neuroepithelial expansion and neuroblast transition. Dev Cell 2013, 27:174–187.

31. Loya CM, McNeill EM, Bao H, Zhang B, Van Vactor D: miR-8 controls synapse structure by repression of the actin regulator enabled. Development 2014, 141:1864–1874.

32. Lu CS, Zhai B, Mauss A, Landgraf M, Gygi S, Van Vactor D: MicroRNA-8 promotes robust motor axon targeting by coordinate regulation of cell adhesion molecules during synapse development. Philos Trans R Soc Lond B Biol Sci 2014, 369.

33. Bolin K, Rachmaninoff N, Moncada K, Pula K, Kennell J, Buttitta L: miR-8 modulates cytoskeletal regulators to influence cell survival and epithelial organization in Drosophila wings. Dev Biol 2016, 412:83–98.

34. Choi IK, Hyun S: Conserved microRNA miR-8 in fat body regulates innate immune homeostasis in Drosophila. Dev Comp Immunol 2012, 37:50–54.

35. Etebari K, Asgari S: Conserved microRNA miR-8 blocks activation of the Toll pathway by upregulating Serpin 27 transcripts. RNA Biol 2013, 10:1356–1364.

36. Kennell JA, Cadigan KM, Shakhmantsir I, Waldron EJ: The microRNA miR-8 is a positive regulator of pigmentation and eclosion in Drosophila. Dev Dyn 2012, 241:161–168.

37. Kennell JA, Gerin I, MacDougald OA, Cadigan KM: The microRNA miR-8 is a conserved negative regulator of Wnt signaling. Proc Natl Acad Sci U S A 2008, 105:15417–15422.

38. Lee GJ, Hyun S: Multiple targets of the microRNA miR-8 contribute to immune homeostasis in Drosophila. Dev Comp Immunol 2014, 45:245–251.

39. Lee GJ, Jun JW, Hyun S: MicroRNA miR-8 regulates multiple growth factor hormones produced from Drosophila fat cells. Insect Mol Biol 2015, 24:311–318.

40. Sander M, Eichenlaub T, Herranz H: Oncogenic cooperation between Yorkie and the conserved microRNA miR-8 in the wing disc of Drosophila. Development 2018, 145.

41. Vallejo DM, Caparros E, Dominguez M: Targeting Notch signalling by the conserved miR-8/200 microRNA family in development and cancer cells. EMBO J 2011, 30:756–769.

42. Zeng C, Fukunaga T, Hamada M: Identification and analysis of ribosome-associated lncRNAs using ribosome profiling data. BMC Genomics 2018, 19:414.

43. Hsu PY, Calviello L, Wu HL, Li FW, Rothfels CJ, Ohler U, Benfey PN: Super-resolution ribosome profiling reveals unannotated translation events in Arabidopsis. Proc Natl Acad Sci U S A 2016, 113:E7126–7135.

44. Ingolia NT, Ghaemmaghami S, Newman JR, Weissman JS: Genome-wide analysis in vivo of translation with nucleotide resolution using ribosome profiling. Science 2009, 324:218–223.

45. Dunn JG, Foo CK, Belletier NG, Gavis ER, Weissman JS: Ribosome profiling reveals pervasive and regulated stop codon readthrough in Drosophila melanogaster. Elife 2013, 2:e01179.

46. Kronja I, Whitfield ZJ, Yuan BB, Dzeyk K, Kirkpatrick J, Krijgsveld J, Orr-Weaver TL: Quantitative proteomics reveals the dynamics of protein changes during Drosophila oocyte maturation and the oocyte-to-embryo transition. Proceedings of the National Academy of Sciences of the United States of America 2014, 111:16023–16028.

47. Enderle D, Beisel C, Stadler MB, Gerstung M, Athri P, Paro R: Polycomb preferentially targets stalled promoters of coding and noncoding transcripts. Genome Res 2011, 21:216–226.

48. Qian J, Zhang Z, Liang J, Ge Q, Duan X, Ma F, Li F: The full-length transcripts and promoter analysis of intergenic microRNAs in Drosophila melanogaster. Genomics 2011, 97:294–303.

49. Lucas KJ, Roy S, Ha J, Gervaise AL, Kokoza VA, Raikhel AS: MicroRNA-8 targets the Wingless signaling pathway in the female mosquito fat body to regulate reproductive processes. Proc Natl Acad Sci U S A 2015, 112:1440–1445.

50. Mackay TFC, Richards S, Stone EA, Barbadilla A, Ayroles JF, Zhu DH, Casillas S, Han Y, Magwire MM, Cridland JM, Richardson MF, Anholt RRH, Barron M, Bess C, Blankenburg KP, Carbone MA, Castellano D, Chaboub L, Duncan L, Harris Z, Javaid M, Jayaseelan JC, Jhangiani SN, Jordan KW, Lara F, Lawrence F, Lee SL, Librado P, Linheiro RS, Lyman RF, et al: The Drosophila melanogaster Genetic Reference Panel. Nature 2012, 482:173–178.

51. Dinger ME, Pang KC, Mercer TR, Mattick JS: Differentiating protein-coding and noncoding RNA: challenges and ambiguities. PLoS Comput Biol 2008, 4:e1000176.

52. Ji Z, Song R, Regev A, Struhl K: Many lncRNAs, 5’UTRs, and pseudogenes are translated and some are likely to express functional proteins. Elife 2015, 4:e08890.

53. Ladoukakis E, Pereira V, Magny EG, Eyre-Walker A, Couso JP: Hundreds of putatively functional small open reading frames in Drosophila. Genome Biol 2011, 12:R118.

54. Pauli A, Valen E, Schier AF: Identifying (non-)coding RNAs and small peptides: challenges and opportunities. Bioessays 2015, 37:103–112.

55. Raj A, Wang SH, Shim H, Harpak A, Li YI, Engelmann B, Stephens M, Gilad Y, Pritchard JK: Thousands of novel translated open reading frames in humans inferred by ribosome footprint profiling. Elife 2016, 5:e13328.

56. Smith JE, Alvarez-Dominguez JR, Kline N, Huynh NJ, Geisler S, Hu W, Coller J, Baker KE: Translation of small open reading frames within unannotated RNA transcripts in Saccharomyces cerevisiae. Cell Rep 2014, 7:1858–1866.

57. Andrews SJ, Rothnagel JA: Emerging evidence for functional peptides encoded by short open reading frames. Nat Rev Genet 2014, 15:193–204.

58. Saghatelian A, Couso JP: Discovery and characterization of smORF-encoded bioactive polypeptides. Nat Chem Biol 2015, 11:909–916.

59. Carlevaro-Fita J, Rahim A, Guigo R, Vardy LA, Johnson R: Cytoplasmic long noncoding RNAs are frequently bound to and degraded at ribosomes in human cells. RNA 2016, 22:867–882.

60. Chugunova A, Loseva E, Mazin P, Mitina A, Navalayeu T, Bilan D, Vishnyakova P, Marey M, Golovina A, Serebryakova M, Pletnev P, Rubtsova M, Mair W, Vanyushkina A, Khaitovich P, Belousov V, Vysokikh M, Sergiev P, Dontsova O: LINC00116 codes for a mitochondrial peptide linking respiration and lipid metabolism. Proc Natl Acad Sci U S A 2019.

61. Khitun A, Ness TJ, Slavoff SA: Small open reading frames and cellular stress responses. Mol Omics 2019.

62. Gault WJ, Olguin P, Weber U, Mlodzik M: Drosophila CK1-gamma, gilgamesh, controls PCP-mediated morphogenesis through regulation of vesicle trafficking. J Cell Biol 2012, 196:605–621.

63. McCarthy DJ, Chen Y, Smyth GK: Differential expression analysis of multifactor RNA-Seq experiments with respect to biological variation. Nucleic Acids Res 2012, 40:4288–4297.

64. Mi H, Muruganujan A, Casagrande JT, Thomas PD: Large-scale gene function analysis with the PANTHER classification system. Nat Protoc 2013, 8:1551–1566.

65. Michel AM, Fox G, A MK, De Bo C, O’Connor PB, Heaphy SM, Mullan JP, Donohue CA, Higgins DG, Baranov PV: GWIPS-viz: development of a ribo-seq genome browser. Nucleic Acids Res 2014, 42:D859–864.

66. Montigny A, Tavormina P, Duboe C, San Clémente H, Aguilar M, Valenti P, Lauressergues D, Combier JP, Plaza S: Data for « Drosophila primary microRNA-8 encodes a microRNA encoded peptide acting in parallel of miR-8 ». available from https://www.ncbi.nlm.nih.gov/Traces/study/?acc=PRJNA645280&o=acc_s%3Aa. 2020.

