## Supplementary Figures for "*Drosophila* primary microRNA-8 encodes a microRNA encoded peptide acting in parallel of *miR-8*"

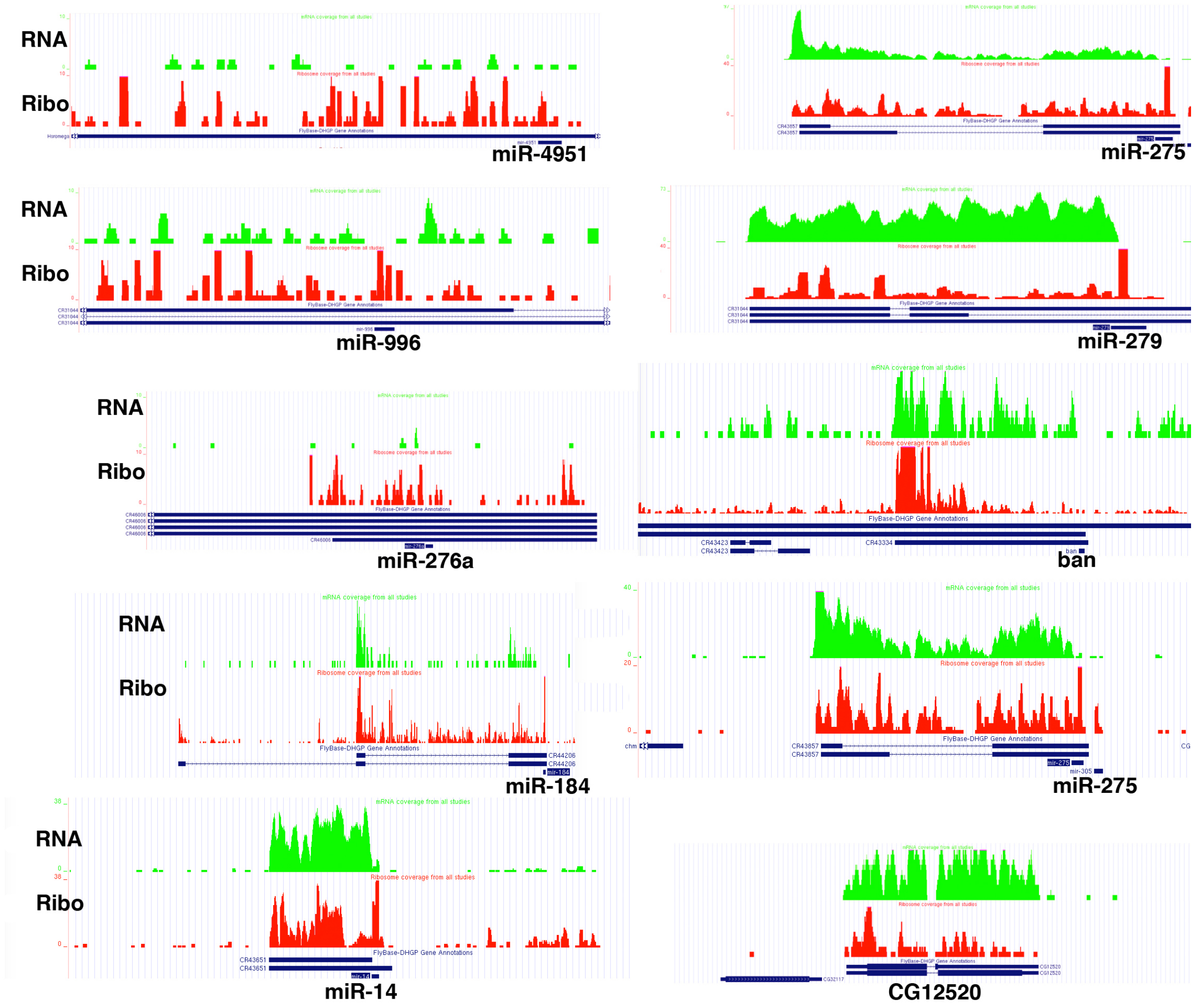


**Figure S1 :** Ribosome covering profiles on several *Drosophila* mi-R genes.
Screen shot of examples of miR genes exhibiting ribosome profiling using the GWIPS-vis browser (https://gwips.ucc.ie) [65]. In green are shown the RNA-seq expression, in red ribosome profiling datas. The coding gene CG12520 is shown for comparison.

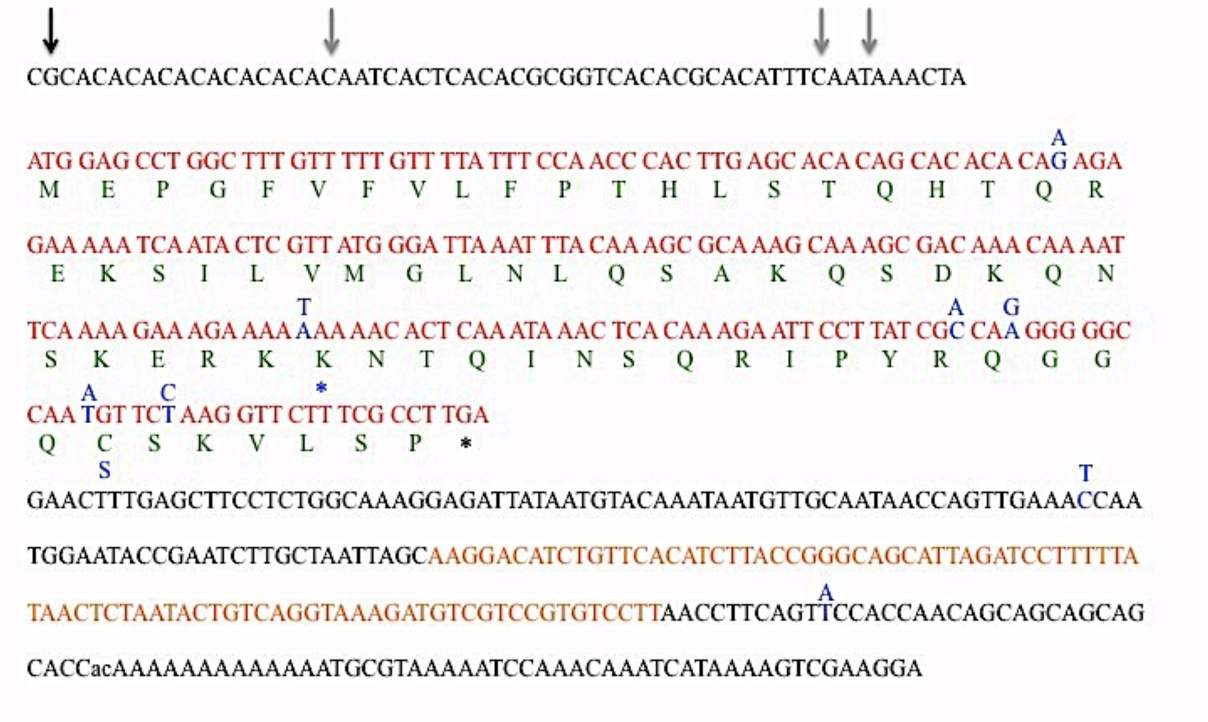


**Figure S2 :** *pri-miR8* sequence.
In red is shown the open reading frame for miPEP-8. In green is listed the miPEP-8 amino acid sequence. In orange is shown the pre-miR-8. In blue are indicated the SNPs detected and the * indicate the premature stop codon.
Arrows indicate the 5' end identified in 5'RACE experiment. Black arrow show the most 5' end identified by 5'RACE and correlating with the 5'ends found in RNA-seq.

A

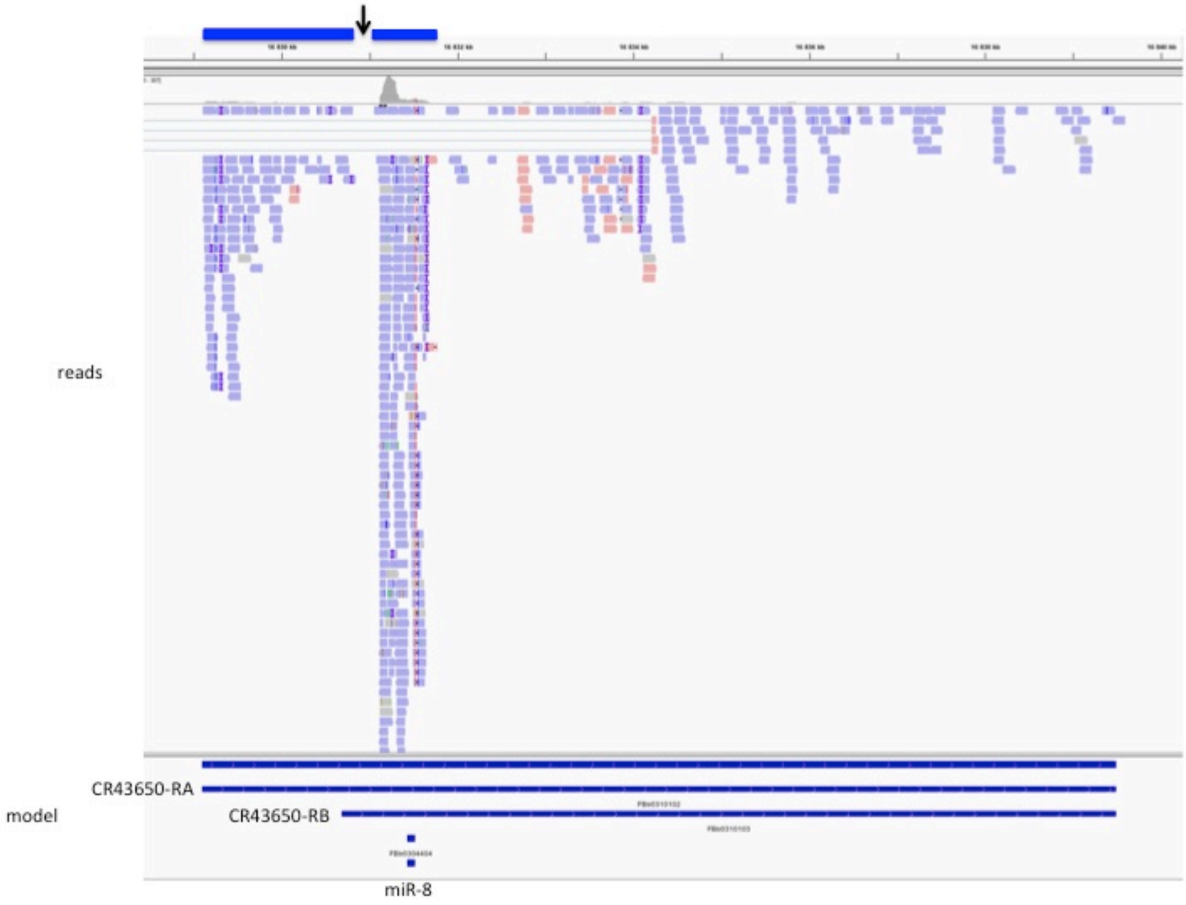


**B**

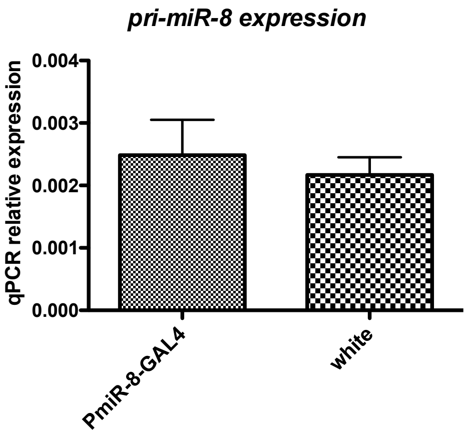


**Figure S3** :
**A** : RNA-seq profil of the *miR-8* locus. Contrasting with the models of long RA and short RB CR43650 transcripts, no covering reads and overlapping reads were detected in the region located upstream the RB transcript suggesting that two different transcriptional units are present (schematized by blue rectangles on top). Arrowhead : Location of the *miR-8* GAL4 line insertion.
**B** : expression of *miR-8* gene in white recipent flies and miR-8 GAL4 flies. Differences are non significant (Mann & Whitney non parametric test). n=6

A

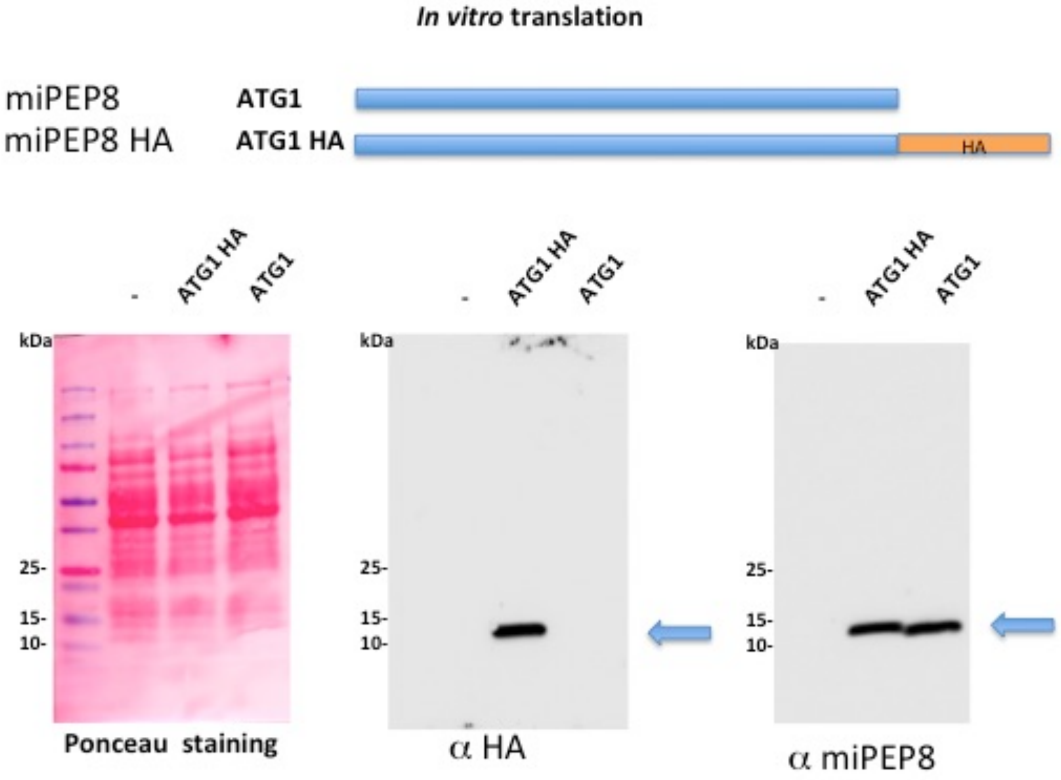

*In vitro* translation
miPEP8
ATG1
miPEP8 HA
ATG1 HA
HA
-
ATG1 HA
ATG1
kDa
25-
15-
10-
Ponceau staining
α HA
α miPEP8


B

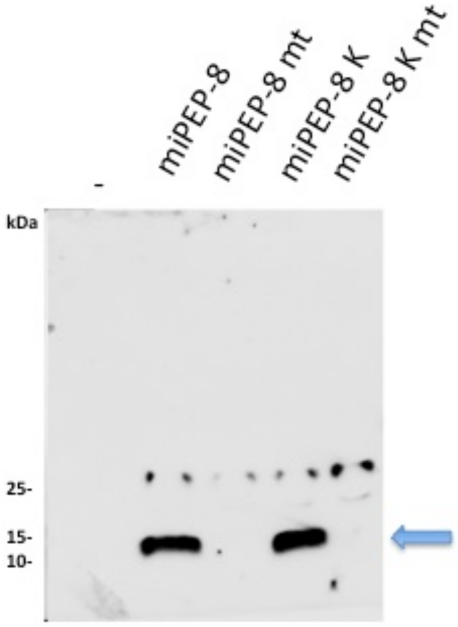

-
miPEP-8
miPEP-8 mt
miPEP-8 K
miPEP-8 K mt
kDa
25-
15-
10-

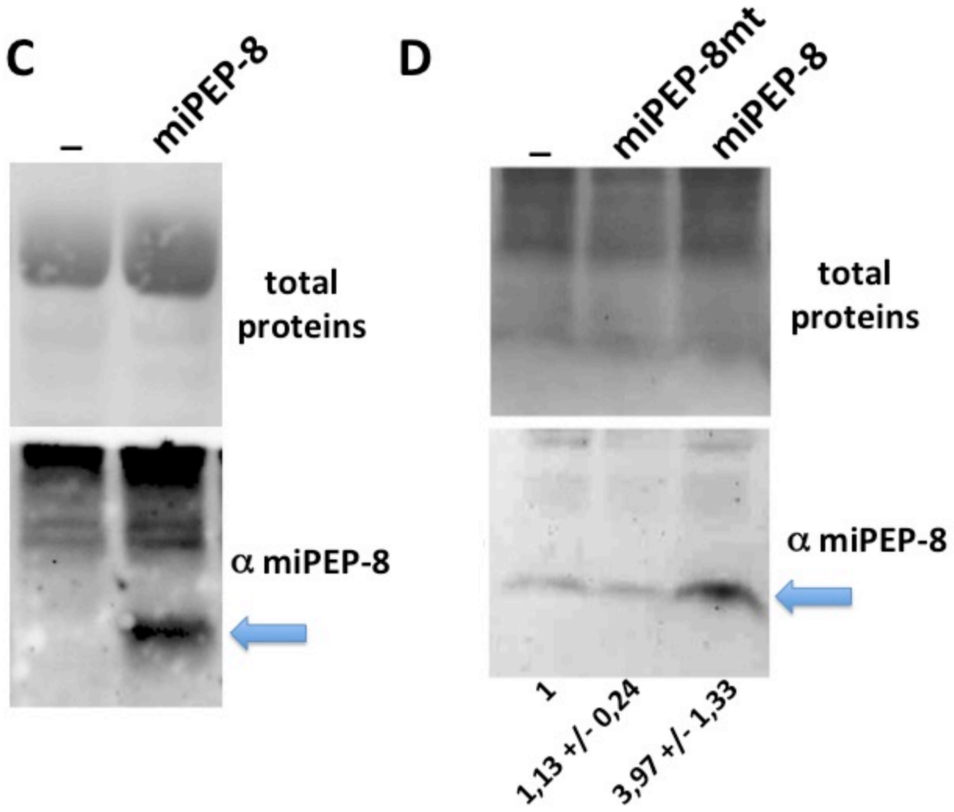


**Figure S4** : anti miPEP-8 antibodies characterization. - indicates programmed lysate with empty vector. **A** : *In vitro* translated miPEP-8 HA were produced in insect cell extracts and subjected to western blot experiments. **B** : the miPEP-8 ORFs ATGs were placed in natural, kozak (K) or mutated (mt) translational context and were tested for their ability to be translated. **C :** *N.benthamiana* agroinfiltrated with pCambia expressing vectors, empty vector (ctrl) or miPEP-8 encoding vector (miPEP-8). Arrows indicate miPEP-8. Note the signal in miPEP-8 overexpressing plants. **D** : miR-8 GAL4 flies were crossed with the respective UAS constructs (ctrl is driver crossed with the recipent white flies). Young Adults (1-2 days old) were subjected to western blot using the purified anti miPEP8 antibody. Arrows indicate miPEP-8. Note the increased signal in miPEP-8 overexpressing flies for the wt construct but not the ATG mutated one.

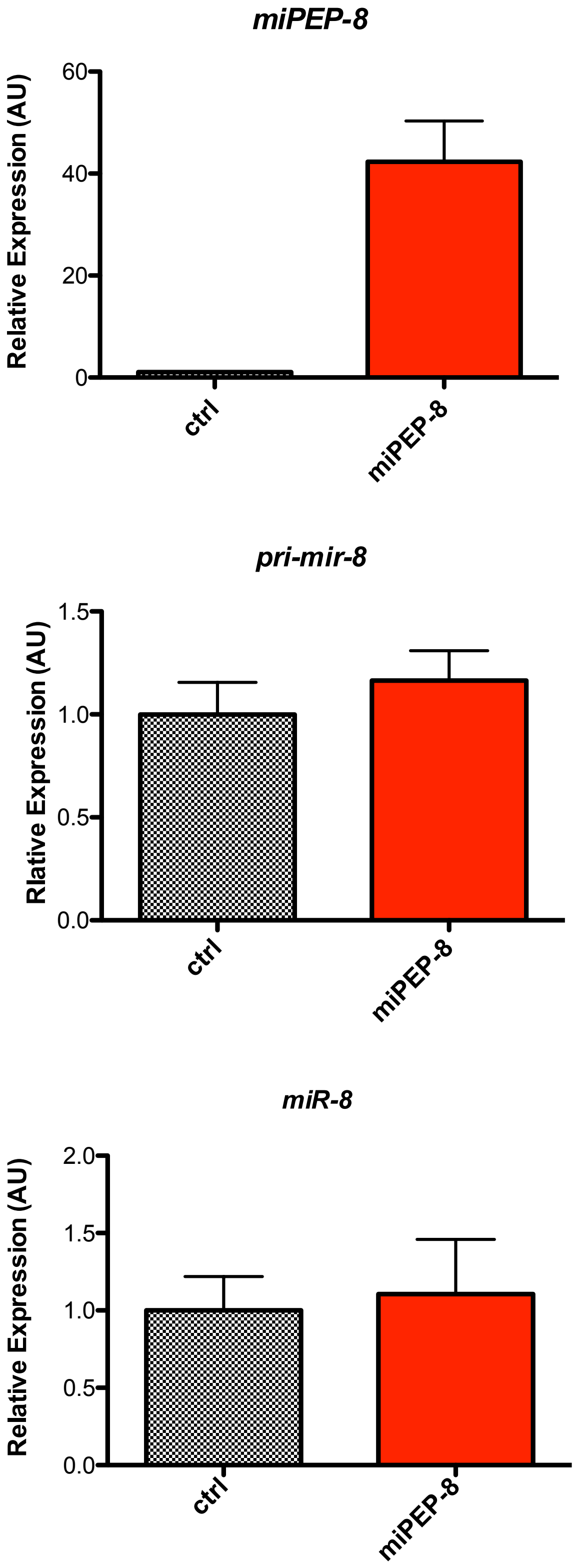


**Figure S5** : qPCR on flies over-expressing miPEP-8 using the *miR-8* GAL4 line as driver. Top panel, the overexpression level is more than 45 fold times higher than the endogenous level (n=9). Middle and bottom panels, endogenous level of *pri-miR-8* (n= 17) and mature miR-8 (n=10) were determined by qPCR. Non significant variation (Mann & Whitney non parametric test) of endogenous *pri-miR-8* or mature *miR-8* was observed upon miPEP-8 overexpression.

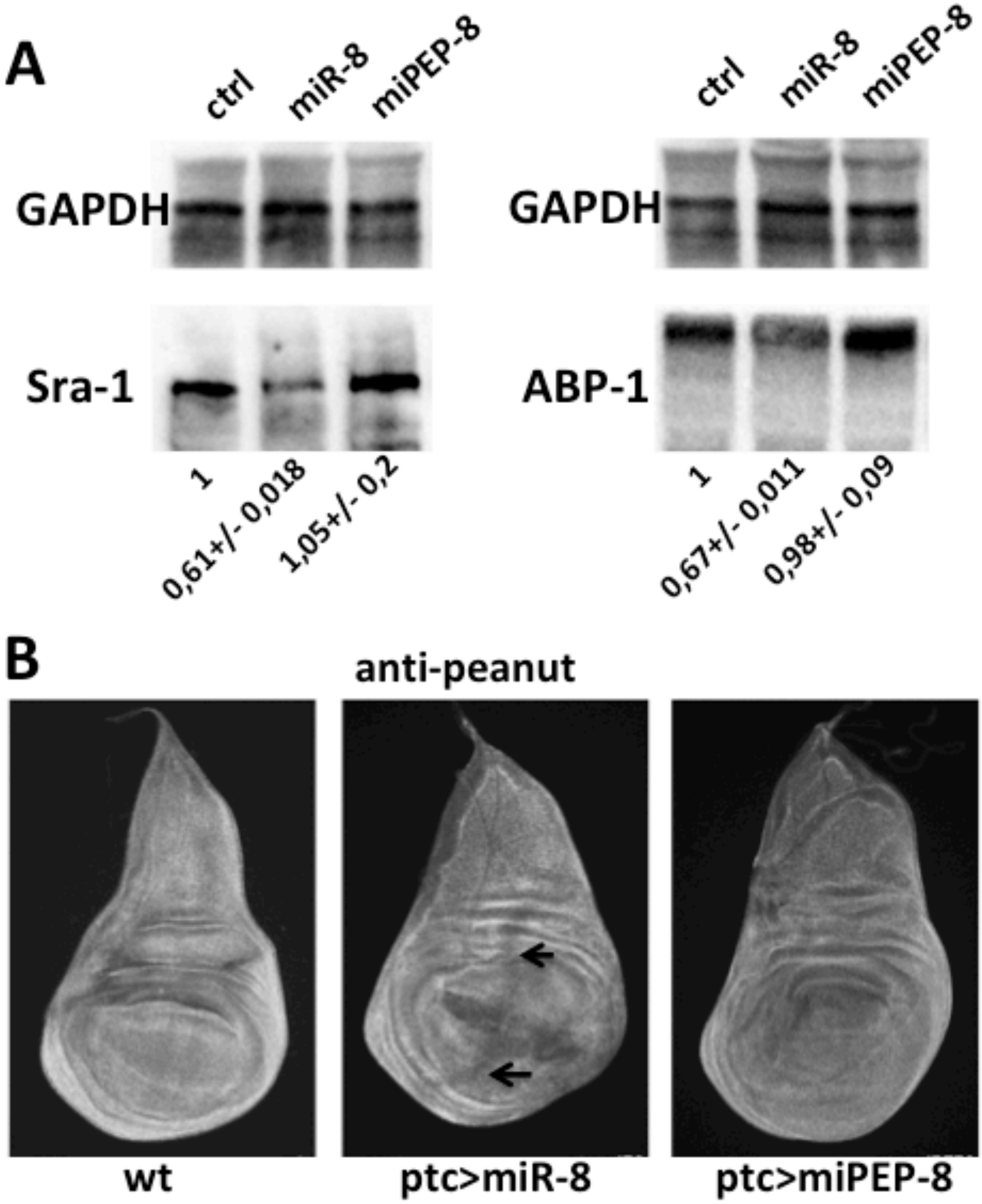


**Figure S6** : Lack of miPEP-8 effect on miR-8 endogenous targets.
**A** : western blot experiment on S2 cells overexpressing *miR-8* or miPEP-8 as indicated. Bottom : average value of the relative expression/GAPDH from two independent experiments with SD indicated. **B** : expression of *miR-8* and miPEP-8 using ptc-GAL4 in wing imaginal discs. Third instard discs are stained with an anti-peanut antibody. Arrows : repression of this Peanut expression in the ptc domain with *miR-8* but not miPEP-8.

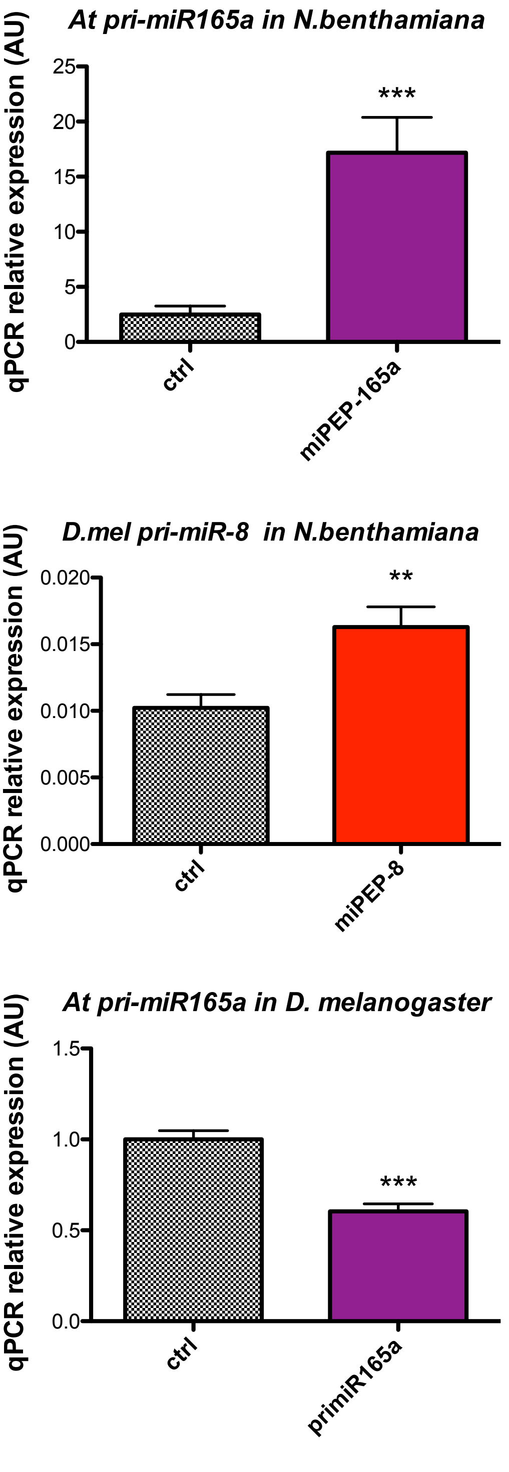


**Figure S7** : qPCR on agroinfiltrated *N.benthamiana* leaves with either *A.thaliana pri-miR165a* (n= 8) or *Drosophila pri-miR-8* (n= 12) expressed from ubiquitous promoter (35S) together with either an empty expression vector (ctrl) or a vector expressing miPEP-165a or miPEP-8 respectively. Bottom transfected *drosophila* S2 cells (n= 13) expressing the *A.thaliana pri-miR165a* together with the p-actin empty vector (ctrl) or a p-actin vector expressing the miPEP-165a. * indicate that the difference is significant with a pvalue<0,01 (**) or 0,001 (***)

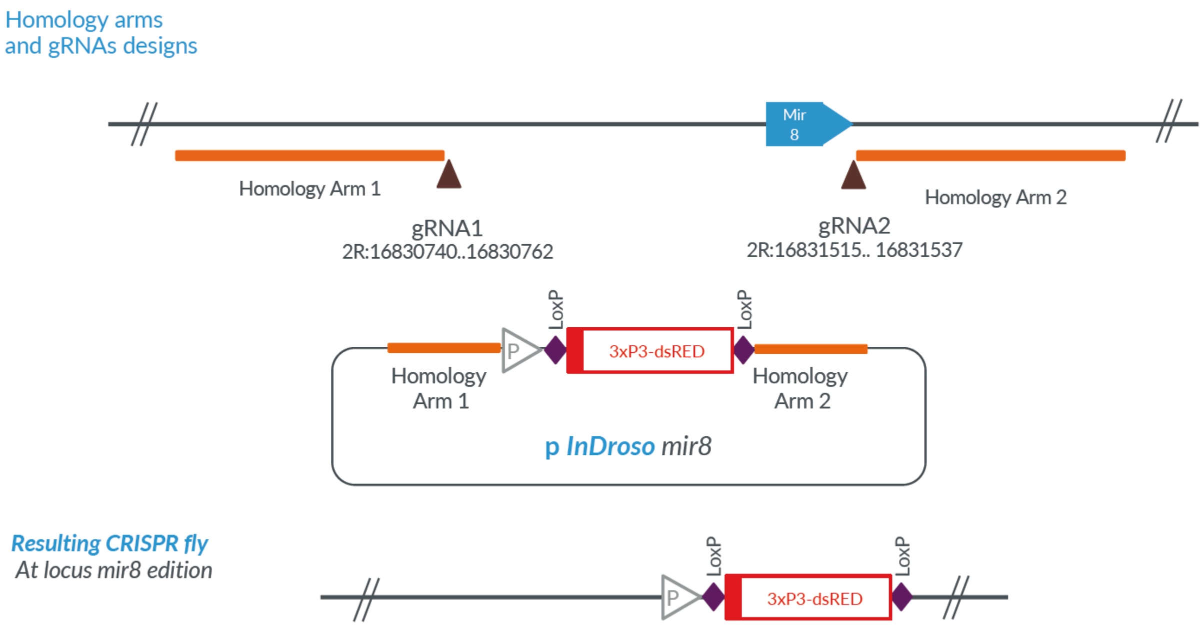


pri-miR-8
Homojogy arm1 - attP site - LoxP – 3xP3 dsRED - LoxP site- Homology arm2
Ccgctg disrupted gRNA1 binding site
Ccttcagtaccaccaac disrupted gRNA2 binding site

//ggtgctcgatcgtgacggtttgctcgcgctctccgctgcgccgctcgtagtgccccaactggggtaacctttgagttctctcagttgggggcgtagataacttcgtataatgtatgctAtacgaagttatCGTA
CGGGATCTAATTCAATTAGAGACTAATTCAATTAGAGCTAATTCAATTAGGATCCAAGCTTATCGATTTCGAACCCTCGACCGCCGGAGTATAAAT
AGAGGCGCTTCGTCTACGGAGCGACAATTCAATTCAAACAAGCAAAGTGAACACGTCGCTAAGCGAAAGCTAAGCAAATAAACAAGCGCAGCTG
AACAAGCTAAACAATCGGCTCGAAGCCGGTCGCCACCATGGCCTCCTCCGAGGACGTCATCAAGGAGTTCATGCGCTTCAAGGTGCGCATGGAG
GGCTCCGTGAACGGCCACGAGTTCGAGATCGAGGGCGAGGGCGAGGGCCGCCCCTACGAGGGCACCCAGACCGCCAAGCTGAAGGTGACCAA
GGGCGGCCCCCTGCCCTTCGCCTGGGACATCCTGTCCCCCCAGTTCCAGTACGGCTCCAAGGTGTACGTGAAGCACCCCGCCGACATCCCCGAC
TACAAGAAGCTGTCCTTCCCCGAGGGCTTCAAGTGGGAGCGCGTGATGAACTTCGAGGACGGCGGCGTGGTGACCGTGACCCAGGACTCCTCC
CTCCAGGACGGCTCCTTCATCTACAAGGTGAAGTTCATCGGCGTGAACTTCCCCTCCGACGGCCCCGTAATGCAGAAGAAGACTATGGGCTGGG
AGGCGTCCACCGAGCGCCTGTACCCCCGCGACGGCGTGCTGAAGGGCGAGATCCACAAGGCCCTGAAGCTGAAGGACGGCGGCCACTACCTGG
TGGAGTTCAAGTCCATCTACATGGCCAAGAAGCCCGTGCAGCTGCCCGGCTACTACTACGTGGACTCCAAGCTGGACATCACCTCCCACAACGA
GGACTACACCATCGTGGAGCAGTACGAGCGCGCCGAGGGCCGCCACCACCTGTTCCTGTAGGGGCCGCGACTCTAGATCATAATCAGCCATACC
ACATTTGTAGAGGTTTTACTTGCTTTAAAAAACCTCCCACACCTCCCCCTGAACCTGAAACATAAAATGAATGCAATTGTTGTTGTTAACTTGTTTA
TTGCAGCTTATAATGGTTACAAATAAAGCAATAGCATCACAAATTTCACAAATAAAGCATTTTTTTCACTGCATTCTAGTTGTGGTTTGTCCAAACT
CATCAATGTATCTTAACCGGTataacttcgtataatgtatgctatacgaagttatccttcagtTccaccaacagcagcagcagcacca----
Caaaaaaaaaaaaatgcgtaaaaatccaaacaaatcataaaa//

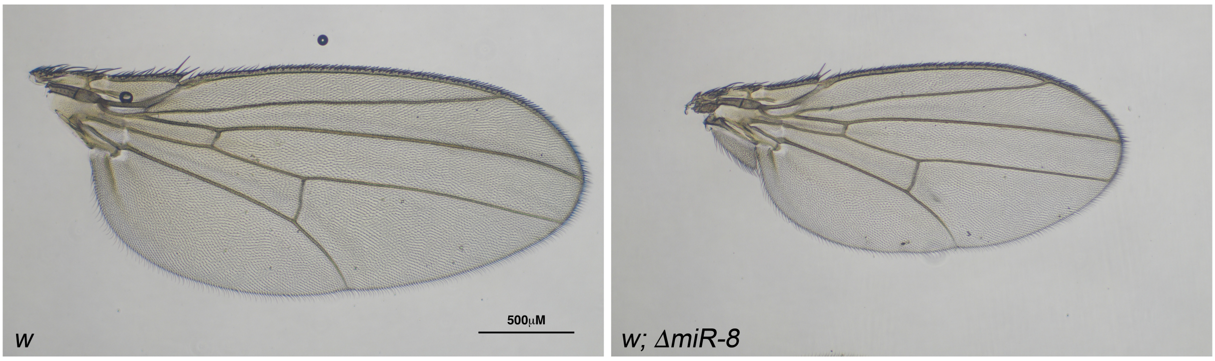


**Figure S8** : scheme of Knock In/ Knock Down strategy ; genomic sequence of the *ΔmiR-8* line and the resulting wing phenotype of the generated *ΔmiR-8* flies

**A**

| gene | symbol | logFC | pvalue (FDR) | updown | reference |
|---|---|---|---|---|---|
| FBgn0015838 | Vang | -0,25 | 0,000100718 | down | Bolin et al, 2016 |
| FBgn0011225 | jar | -0,41 | 3,07216E-11 | down | Bolin et al, 2016 |
| FBgn0005672 | spi | -0,34 | 4,63246E-09 | down | Morante et al, 2013 |
| FBgn0036372 | Abp1 | -0,18 | 0,01777008 | down | Bolin et al, 2016 |
| FBgn0262716 | Arp3 | -0,22 | 0,002244126 | down | Bolin et al, 2016 |
| FBgn0013726 | pnut | -0,25 | 6,25716E-05 | down | Bolin et al, 2016; Eichenlaub et al, 2016 |
| FBgn0003514 | sqh | -0,26 | 9,0503E-05 | down | Bolin et al, 2016 |
| FBgn0038320 | Sra-1 | -0,28 | 2,44587E-05 | down | Bolin et al, 2016 |
| FBgn0034970 | yki | -0,32 | 1,82097E-06 | down | Umegawachiet al, 2017; Sander et al, 2018 |
| FBgn0001257 | ImpL2 | -0,35 | 6,13257E-06 | down | Lee et al, 2015 |
| FBgn0036141 | wls | -0,36 | 1,78976E-07 | down | Kennel et al, 2008 |
| FBgn0034709 | Swim | 0,82 | 3,1208E-15 | up | Lucas et al, 2015 |
| FBgn0034407 | DptB | 1,00 | 0,000544687 | up | Choi & Hyun, 2012 |

**B**

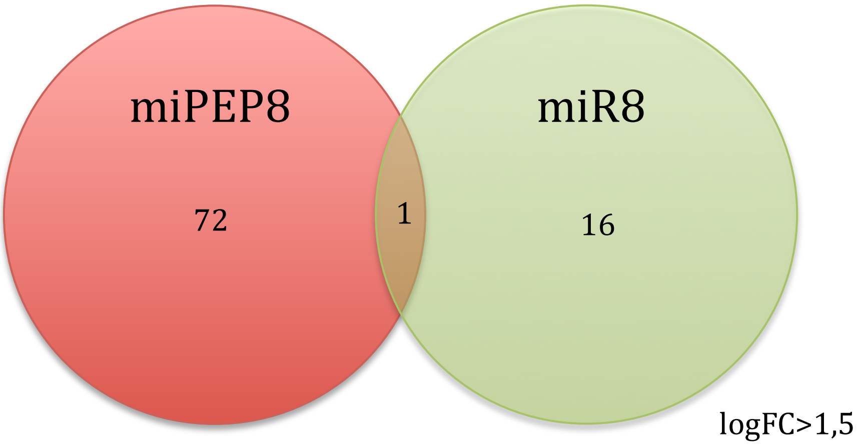


**Figure S9** : miPEP-8 and *miR-8* deregulated genes in S2 cells overexpressing miPEP-8 or miR-8. **A** : published *miR-8* targets identified in this study that are highly significant (see FDR value) with a logFC > 0,25. **B** : venn diagram for significant deregulated genes (number indicated in the center of each circle) with a logFC > 1 ,5.

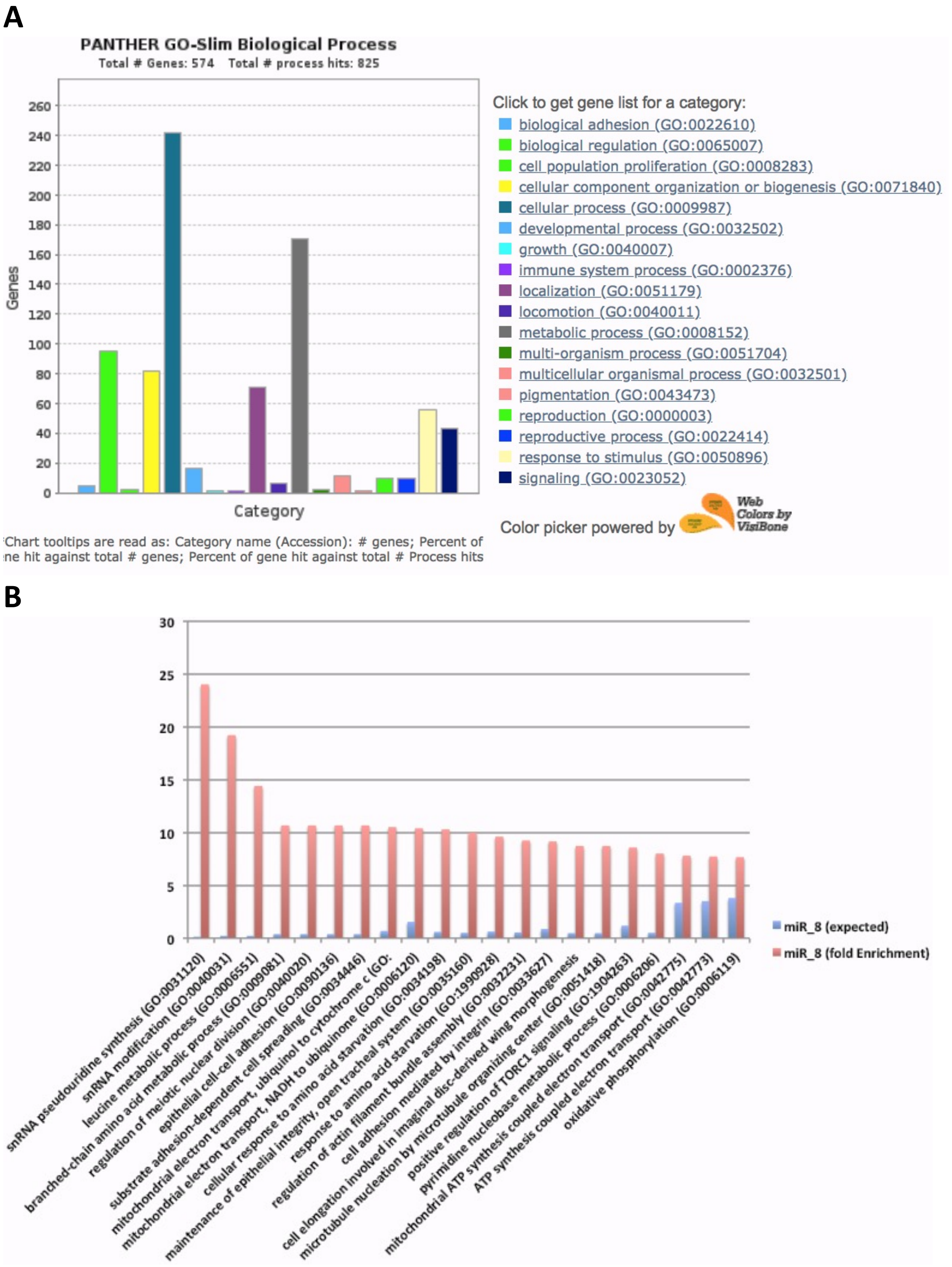


**Figure S10** : GO of transcriptome of *miR-8* specific genes. **A** : Most significant regulated biological processes. **B** : Fold enrichment of the most representative biological activities controled by *miR-8*. Note enrichment of epithelial cell adhesion, and integrity, regulation of actin filament assembly, wing imaginal disc morphogenesis.

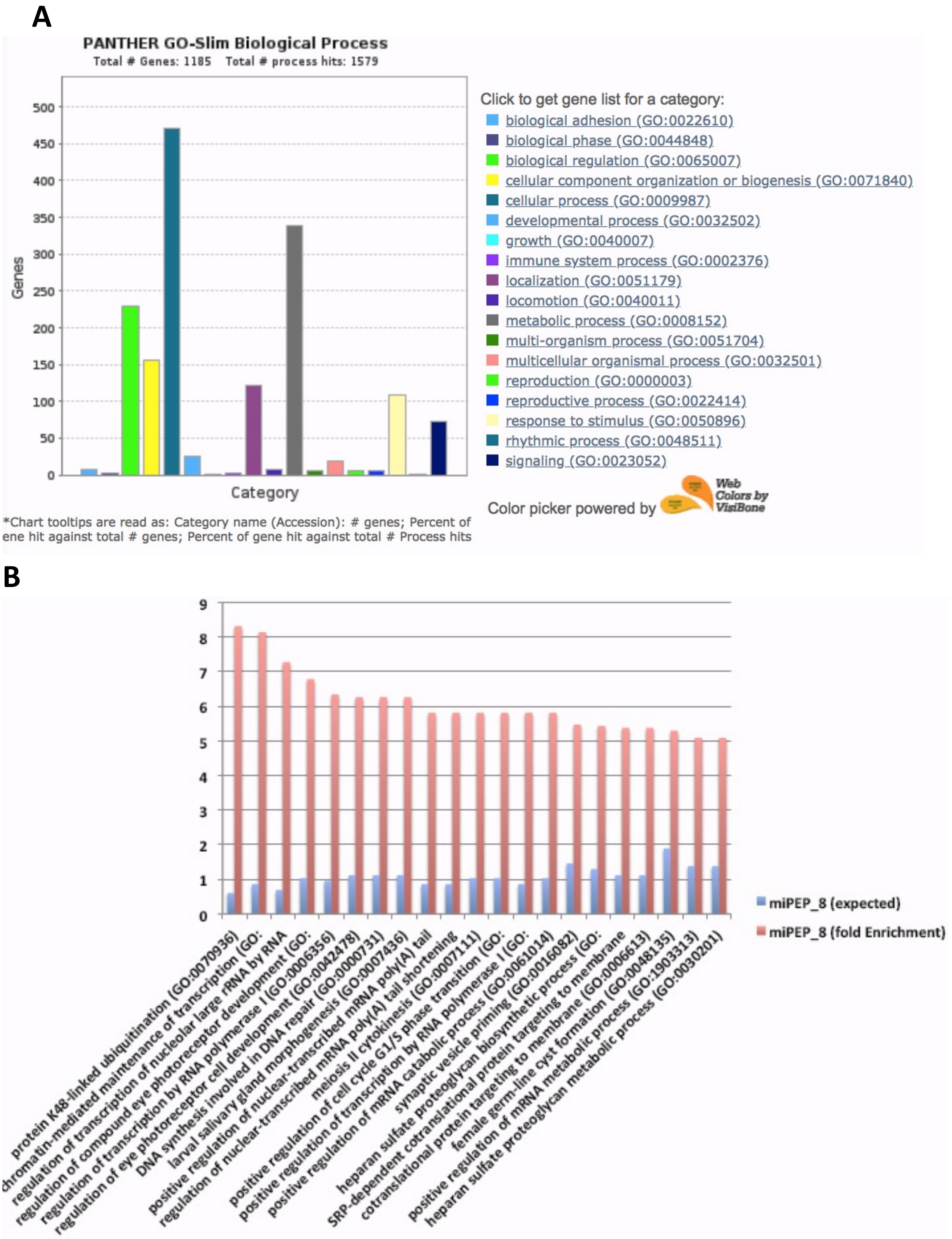


**Figure S11** : Gene Ontology of transcriptome of miPEP-8 specific genes. **A** : Most significant regulated biological processes. **B** : Fold enrichment of the most representative biological activities.

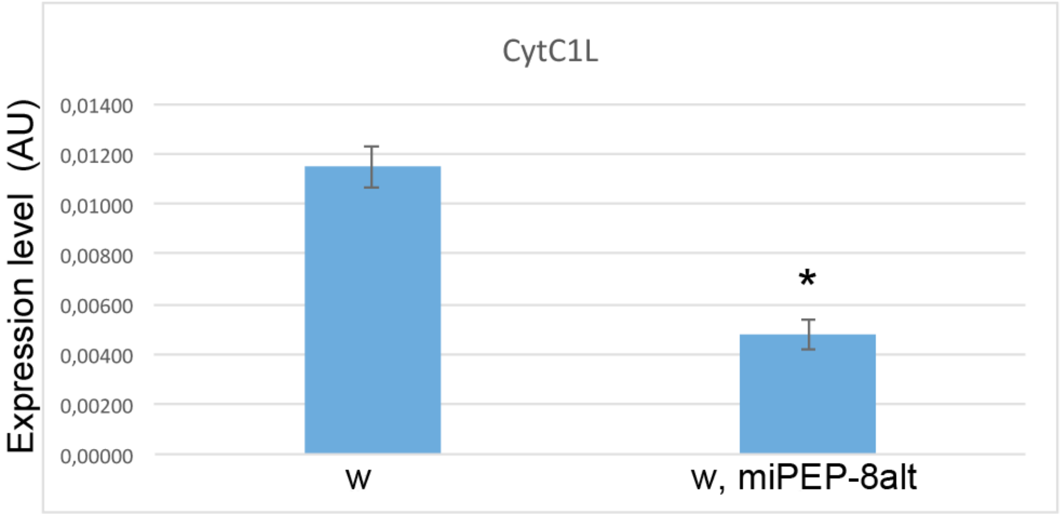


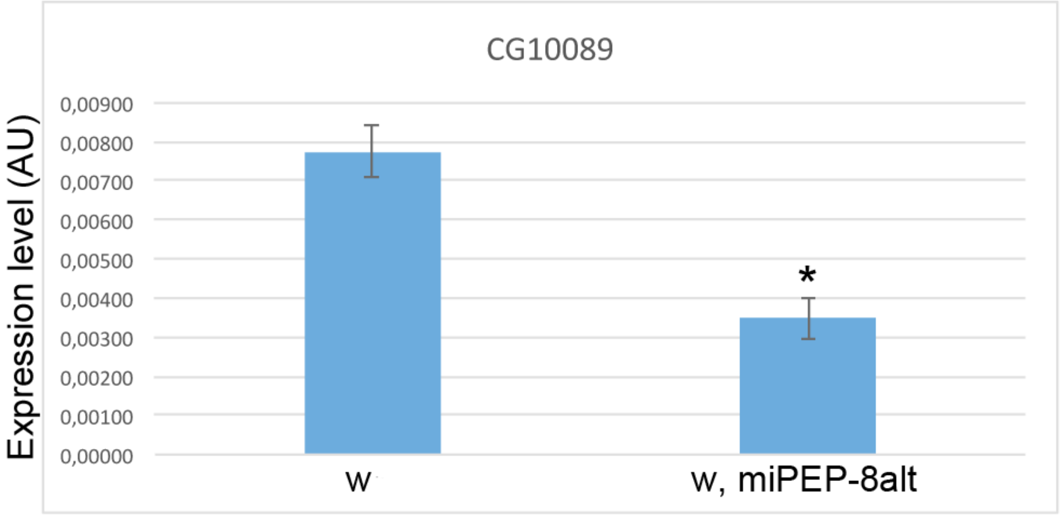


**Figure S12** : Modulation of two miPEP-8 regulated genes analysed by qPCR in white (wt miPEP-8) and in w ; miPEP-8 mutated adult flies (miPEP-8alt) (N= 6 and 8 respectively). The differences are significant (p<0,05).

| | |
|---|---|
| AGATCGTGAAGAAGCGCACCAAGC | RP49 q5 |
| GCACCAGGAACTTCTTGAATCCGG | RP49 q3 |
| CGAGACCTACTGCATCGACA | Dm tub q5 |
| AGGTCACCGTATGTGGGTGT | Dm tub q3 |
| AGGACACGGACGACATCTTT | premiR8 q3 |
| TCTTACCGGGCAGCATTAGA | premiR8 q5 |
| TCGCCTTGAGAACTTTGAGC | primiR8 q5 |
| TGATTTGTTTGGATTTTTACGC | primiR8 q3OK |
| TTGTTTGTCGCTTTGCTTTG | primiR8 q3 quatuor |
| GAGCCTGGCTTTGTTTTTGT | PrimiR8q5quatuor |
| GGGGGTAATACTGTCAGGTAAA | New mir8fwqRT |
| CCAGTGCAGGGTCCGAGGTA | New universal rv stemloop |
| GCTTCGGCTTAATGATGGTC | U14fwqRT |
| GGGTGTGATTCTGCTTGTCA | snoR442fwqRT |
| GTCGTATCCAGTGCAGGGTCCGAGGTATTCGCACTGGATACGACAGTCAG | stemloopRTU14 |
| GTCGTATCCAGTGCAGGGTCCGAGGTATTCGCACTGGATACGACGATCAG | stemloopRTsnoR442 |
| GTCGTATCCAGTGCAGGGTCCGAGGTATTCGCACTGGATACGACGACATC | SL RT miR8 |
| CTTCAGAACCGGAGACCGAC | Cyt C1L Fw |
| TCTTGCGAGACTTGAGCGTT | Cyt C1L Rv |
| GGAGCAACTGGATCGCACTA | CG10089 Fw |
| GCAGTTACCCTCGCAGATGT | CG10089 Rv |

**Figure S13** : sequences and list of primers used in real time quantitative PCR experiments.
